# Frontopolar cortex stimulation induces prolonged disruption to counterfactual processing: insights from altered local field potentials

**DOI:** 10.1101/2024.11.26.625398

**Authors:** Matthew Ainsworth, Juan M. Galeazzi, Carlos Pedreira, Mark G. Stokes, Mark J. Buckley

## Abstract

Frontopolar cortex (FPC) is a large, anterior sub-region of prefrontal cortex found in both humans and non-human primates (NHPs) and is thought to support monitoring the value of switching between alternative goals. However, the neuronal mechanisms underlying this function are unclear. Here we used multielectrode arrays to record the local field potentials (LFPs) in the FPC of two macaques performing a Wisconsin Card Sorting Test analogue and found that bursts of gamma and beta in FPC tracked counterfactual not current rule value. Moreover, we show that brief high-frequency microstimulation to a single trial causally affects both LFP activity in FPC, as well as rule-guided decision-making across successive trials. Following stimulation of FPC we observed reduced exploration of the counterfactual rule pre-rule-change, as well as a delayed adaptation to the newly relevant following post-rule-change. A similar, multi-trial time-course disturbance to beta and gamma activity within FPC was also induced following single-trial microstimulation. These findings conclusively link neuronal activity in FPC with behavioural monitoring of the value of counterfactual rules and provide neural mechanistic insights into how FPC supports rule-based decision-making.

**Significance statement:** Increasing evidence from human and non-human primates has prompted theories linking FPC with the control of exploration during decision-making. However, it is current unclear how the neuronal activity within frontal pole supports exploratory decision-making processes. Here we show that rhythmic activity, in the beta and gamma bands, recorded from FPC is correlated both with outcome of the previous choice, and the value of switching to an alternative choice. Furthermore, we show that disrupting beta and gamma activity within FPC causally influences exploratory decision-making: initially decreasing exploration before impairing adaptation to abstract rule changes. Together these findings provide the first mechanistic insight into how the neuronal activity within FPC can support exploratory behaviour.

## Introduction

Frontopolar cortex (occupied by area 10) is one of the larger sub-regions of primate prefrontal cortex [1] but our understanding of its contributions to cognition remains nascent. Despite often being regarded as an area supporting some of the highest-order cognitive processes in humans including reasoning, planning, and abstraction, no clear consensus has emerged as to its precise role or mechanistic contribution. One recent proposal that emerged from considering both the human and non-human primate (NHP) literature is that FPC influences cognitive flexibility, maintaining an appropriate balance between exploration and exploitation, evaluating the relative value of alternatives, and monitoring competing goals [2]. This contribution of primate FPC to cognition may be distinct from that of other posterior prefrontal regions as supported by a number of correlative findings from both human neuroimaging [3–6] as well as causal findings from circumscribed FPC lesion behavioural studies in macaque [7–10]. Damage to FPC decreases behavioural flexibility in both humans and NHPs. Patients whose lesions encroach substantially upon FPC are more impaired at multi-tasking [11], task-switching [12], and prospective memory-based task-selection [13], than patients whose lesions do not encroach so extensively upon FPC. Moreover, the aforementioned exploit/explore trade-off can also explain observations of (otherwise counterintuitive) FPC lesion-induced performance enhancements in certain situations. For example, patients whose rostral prefrontal lesions encompassed FPC outperformed non-lesioned controls when required to keep exploiting the previous rule [12]. Similarly, macaques with circumscribed FPC lesions remained well focused on a complex primary task and continued to effectively exploit rewards therein despite distractions, significantly outperforming normal controls with intact FPC whose performance dropped to chance with the same distractions [6].

Nonetheless, our understanding of the specific contribution of FPC to cognition is stymied by a paucity of neuronal data [14]. There are no published reports of neuronal activity from human FPC, and only limited recordings from macaque FPC. Initial studies highlighted involvement of FPC in monitoring self-generated decisions [15–17]. More recent electrophysiological recordings from FPC have demonstrated neurons which encode signals related to the monitoring of actions of other agents [18] or goals during fast learning [19]. Both findings are consistent with previous literature: several fMRI studies have shown activation of FPC when monitoring social-interactions [20, 21] and lesion of FPC has been shown to cause deficits in fast learning [8]. However, whilst these findings are consistent with the purported role for FPC in monitoring the relative benefits of exploitation versus exploration [2] the nature of the neuronal mechanisms in FPC that support these explore/exploit processes in general remains unclear. For example, nothing is known of the frequency-specific oscillatory dynamics within FPC, and the implications this activity has for information encoding, nor whether disruption of such activity in turn influences this brain region’s contribution to monitoring counterfactuals/alternatives.

In this study we recorded the local field potentials (LFPs) from FPC of two monkeys (*Macaca mulatta*) whilst they performed an analogue of the Wisconsin Card Sort Task (WCST), a well-established behavioural task that involves abstract rule-guided decision-making and periodic reversals between choices based on the changing values of alternative uncued abstract rules (i.e. matching-by-colour, and matching-by-shape) [7, 22]. Using a combination of reinforcement learning (RL) modelling, recordings from arrays in FPC, and direct electrical micro-stimulation through electrodes in these arrays, we tested the causal contribution of FPC neuronal activity to rule-guided (and counterfactual rule-guided) decision-making.

We present mechanistic evidence linking beta and gamma frequency activity with counterfactual rule valuation, and exploratory behaviour. We show that gamma frequency activity within FPC, aligned to the animals’ choices, tracked reward/feedback, whilst bursts of gamma and beta activity in FPC tracked the value of the counterfactual (to the currently reinforced rule). Furthermore, targeted causal manipulation of endogenous neuronal activity within FPC using electrical microstimulation significantly modified the amplitude of the LFP recorded from FPC as well as the animal’s performance in the task. Specifically, stimulation of FPC to a single trial about 10 trials before the next rule-change both decreased the exploration of the counterfactual rule during trials leading up to the uncued rule/block change, and also increased the animal’s perseverance after the rule change. These changes were accompanied by trial specific changes in beta and gamma activity in FPC. Together these findings provide further evidence linking FPC with control of exploratory behaviour and provides the demonstration of a mechanistic link between LFP activity recorded from FPC and variables needed to guide exploratory behaviour.

## Results

### Animals’ behavioural choices are influenced by the value of the counterfactual rule

On each trial of the WCST analogue the monkeys were required to select one of three choice stimuli, presented around a central sample stimulus, based on either the colour or shape of that sample stimulus (Fig 1A and Materials and Methods for further details). Of these three choice stimuli one matched the sample in colour, one in shape, whilst the third, a distractor stimulus, matched none of the features of the sample. The correct rule to apply to gain reward on any given trial (i.e. to colour-match or shape-match) was never cued to the animal and so had to be learnt by trial-and-error and then retained in memory across trials. The reinforced rule alternated without announcement between blocks whenever monkeys attained a performance criterion of 85% correct in 20 consecutive trials.

**Fig 1.**
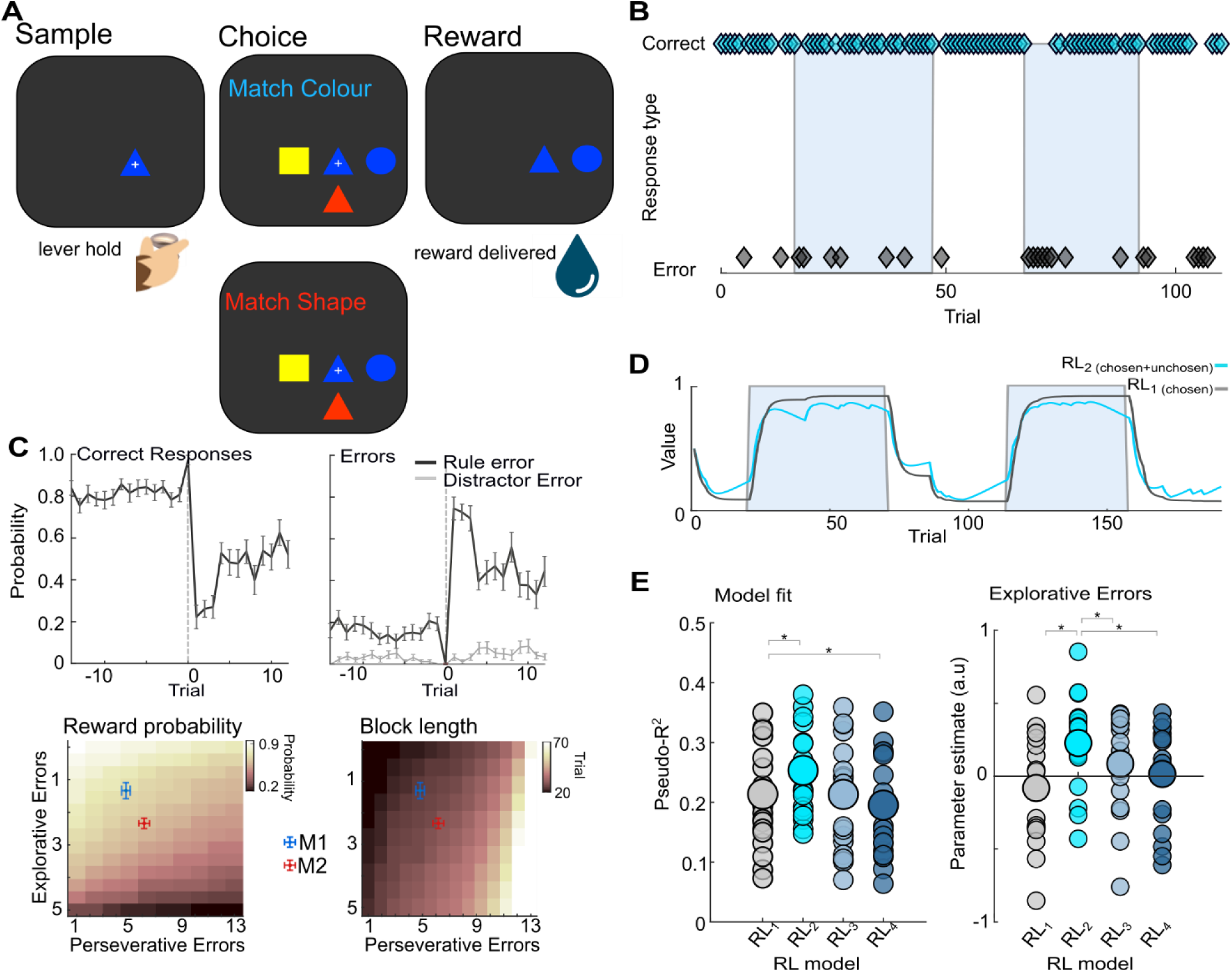
Reinforcement learning modelling of monkeys’ behavioural performance of the WCST analogue. **A.** Task schematic: monkeys matched a central sample stimulus-item to one of three surrounding choice-items based either on a colour or shape-matching rule; the currently correct rule was uncued and changed on a block-by-block basis whenever they attained 85% correct in 20 consecutive trials. **B.** Behavioural performance during a single recording session. Trial outcome indicated with cyan (correct response, top) or black (rule errors, bottom) diamonds. Shape-matching block indicated by blue background and colour-matching block by white. Note rule errors occur throughout blocks. **C.** *Upper:* averaged performance of both animals, aligned to block changes (trial 0). Mean correct responses (*left*), and rule and distractor errors (*right*) shown ±SEM. Note consistent explorative rule errors prior to block change. *Lower*: Colourmaps showing the idealised expected reward (*left*) and block length (*right*) relative to the number of explorative and perseverative errors. Cross hairs show the animals actual performance (mean ±SE). **D.** The predicted value of the shape rule from a single session, obtained from two RL models of interest. In the *RL_1chosen_* model *(black*) only the value of the chosen rule is updated, in the *RL_2chosen+unchosen_* model *(cyan*) both values are updated on every trial. Note fluctuations in predicted value, corresponding to rules errors made by animals, obtained with the *RL_2chosen+unchosen_* model. **E.** Model fit of defined as Mcfaddens Psuedo r^2^ for four models (*left*) *RL_1chosen_* (*gray*) and *RL_2chosen+unchosen_* (*cyan*) as well as two control models (*RL_3_* & *RL_4_, see Materials and Methods for details*). GLMs (*right*) comparing explorative errors predicted by all four models (*RL_1-4_*) with those made by M1 and M2 during the last 7 trials of a block. Analysis revealed *RL_2chosen+unchosen_* provided a significantly better prediction of explorative rules errors at the end of a rule block and a better overall prediction of the animals’ choices. Individual sessions indicated by single dots. Large circles denote the mean for each model.

Both monkeys received extensive training on this task prior to this study. For recording-only sessions collected in this study animals completed an average of 5 blocks per session (mean block/rule changes per session were 6.25 and 3.5 for M1 and M2 respectively). On average there were 127 correct trials per recording session (151 and 102 correct trials/session for M1 and M2 respectively). Visualisation of the correct and incorrect responses made by monkeys (Fig. 1B & C) revealed several notable aspects of the animals’ behaviour. The majority of incorrect responses made by both monkeys were “rule errors” in which the monkeys chose a stimulus corresponding to the incorrect/counterfactual rule (i.e., matching the shape rather than the colour of a stimulus, or vice-versa). By contrast the monkeys rarely chose the distractor stimulus which neither matched the sample in colour nor shape (Fig 1C) indicating that on the vast majority of error trials they still know a matching rule applies even if they do not always implement the correct one.

The frequency with which the monkeys made rule errors varied markedly relative to rule changes. During the initial trials on each block (immediately after rule changes), monkeys made numerous rule errors as they perseverated with the previous rule (‘perseverative errors’). However, the monkeys continued to make some rule errors throughout the block, including on the trials before a rule-switch when the correct rule to exploit was well established (‘explorative errors’, Fig 1B & C). These explorative errors were made with comparable reaction times to correct choices (Fig S2), suggesting that the animals planned such exploration early in trials (and hence avoided any potential reaction time penalty associated with switching [23]. Comparison of errors made by both animals with idealised models predicting reward delivery and block length (Fig 1C) relative to rule errors (both explorative and perseverative) revealed that neither animal used the optimum behavioural strategy (i.e., determining the current rule and exploiting it unwaveringly until the rule changes) to perform the task. Instead, both animals made a small number of explorative errors throughout testing sessions which avoided causing a significant behavioural penalty (either in terms of reward receipt or increased block length). These data suggest that both animals employed an alternative behavioural strategy in which they periodically tested the counterfactual rule throughout the block. Crucially, this strategy attaches a value to the counterfactual (unchosen) rule, suggesting that the monkeys simultaneously maintained and updated the value for both the current (i.e. correct) and the counterfactual (i.e. incorrect) rule, with both updated in response to the trial outcome.

To confirm whether this alternative strategy, requiring estimated values for both rules, could account for the monkeys’ behaviour we modelled the value associated with each rule by fitting a number of RL models [24, 25] to the choices made during each session (Fig 1D & Fig S1). In the first model (RL _1 chosen_), only the value associated with the chosen rule was updated (on a trial-by-trial basis), with the value of the unchosen rule maintained until selected again. The second model (RL _2 chosen+unchosen_) differed in that the values associated with both the chosen and unchosen rules were updated on every trial (note each rule was updated in the same direction, but with differing values for each learning rate, see Materials and Methods for details). In addition, we fit two further RL models to exclude the possibility that these explorative errors arose because animals were increasingly unable to remember the correct rule to exploit throughout blocks, or because they were momentarily distracted and chose incorrectly (Fig S1, RL _3 random noise_ and RL _4 point disruption_ respectively).

Examination of the goodness-of-fit achieved by all four RL model variants, by means of a repeated measure ANOVA (Fig1E, one between subject factor, Monkey, with two levels M1 and M2, and one within subject factor, Model, with four levels RL _1-4_, see Materials and Methods for details), revealed a significant difference between the four models (F_(3,38)_ = 35.61, p = 2.93×10^−12^). The fit achieved by updating the value of both the chosen and unchosen rules for each trial was significantly better than that from updating only the value of the chosen rule (Fig 1E, Pseudo r^2^ for RL _2 unchosen + chosen_ vs. RL _1 chosen_, 0.25 ±0.009 vs. 0.21 ±0.011 respectively, Cohen’s d =1.59, p = 5.54×10^−5^, post-hoc t-test with Bonferroni correction for multiple comparisons). By contrast, the fit achieved by including a random noise component (representing an inability to recall the correct rule), did not improve the model fit (Pseudo r^2^ for RL_3 random noise_ 0.21 ±0.026, Cohen’s D, 0.05 p = 1.00), whilst the inclusion of a function capturing single trial distractions within the model resulted in significantly worse fit than the original model (Pseudo R^2^ for RL_4 point disruption_, 0.19 ±0.012, Cohen’s D, 0.88, p = 0.013). Examination of model fit in both animals separately confirmed RL_2 chosen + unchosen_ provided a significantly better fitting model in both animals (Fig S3)

Session-by-session analysis of the explorative rule errors predicted by each RL model against the explorative errors made by the monkeys at the end of each block (Fig 1E) confirmed that the increase goodness of fit achieved by considering both the current and counterfactual rule (RL _2 chosen + unchosen)_ was due to improved prediction of explorative errors made by both animals. A further repeated measures ANOVA (one between subject factor, Monkey, with two levels M1 and M2, and one within subject factor, Model, with four levels RL _1-4,_ see Materials and Methods for further details) revealed that prediction of explorative errors differed significantly depending on RL model (F_(1,16)_ = 10.47, p = 5.18×10^−3^), and that the RL _2 chosen+ unchosen_ provided a significantly better model of explorative rule errors made at the end of a block than all other models.

### Local field potential activity in FPC correlates with the counterfactual rule value and reward

To determine the functional relevance of the LFP activity present in FPC (array locations shown in Fig 2A & Fig S4) whilst monkeys performed the WCST, we first calculated spectrograms with alignments to key task-epoch onsets, namely on-screen presentation of the sample and the targets, as well as the moment monkeys made a choice to the screen which initiated feedback. These analyses revealed a clear and robust response in the high gamma frequency band (55-95 Hz) which peaked 200-220ms after the animals expressed their choice (Fig 2B). By contrast, there was limited evidence of gamma frequency activity aligned to the presentation of either the sample or targets (Fig S5). Further analysis, calculated on an electrode-by-electrode basis, revealed that gamma activity aligned to the choice was widespread across the FPC-arrays in both animals (Fig 2B). The majority of electrodes in M1 (25 of 32 electrodes) and just under half in M2 (26 of 64 electrodes) exhibited significant gamma frequency responses without excessive 50 Hz mains noise (significant at p=0.01, independent t-test, not cluster corrected, see Materials and Methods for details). Comparison of choice aligned activity recorded from both monkeys revealed no significant difference in the peak frequencies observed in both animals (Fig S6, repeated measure ANOVA Monkey by Frequency interaction, f_(2,26)_=3.31, p=0.067).

**Fig 2.**
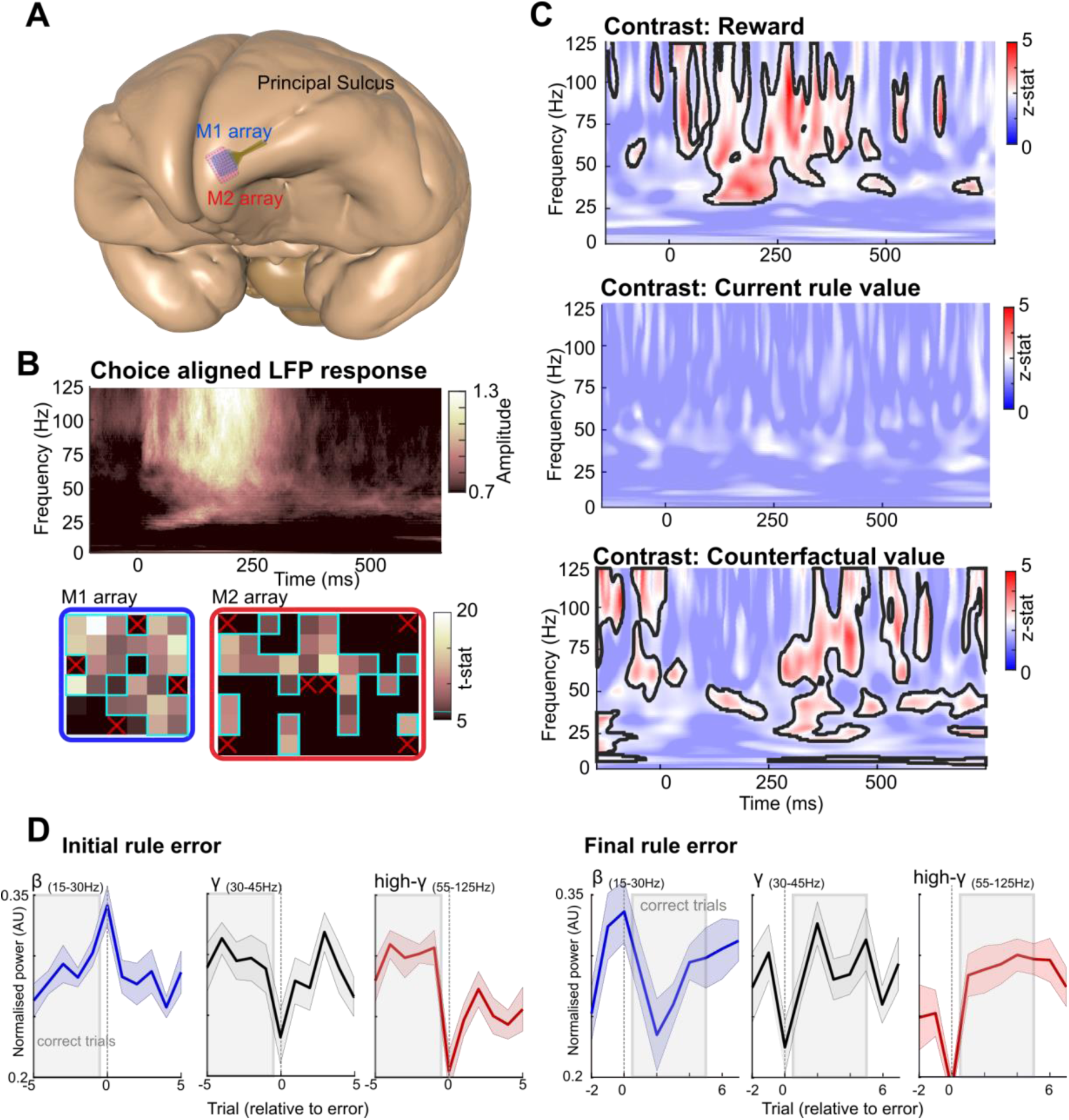
LFP activity recorded from FPC tracks the predicted value of the counterfactual rule. **A.** Illustration showing the location of Utah arrays implanted into FPC of M1 (32 channels, blue) and M2 (64 channels, red); post-study analysis confirmed both arrays implanted in the most rostro-lateral portion of FPC, with the entire array located rostral to the anterior tip of Principal Sulcus. **B.** Choice aligned spectrogram showing a strong gamma frequency response (*top)*. Array maps showing choice responses across the arrays implanted in M1 (blue) & M2 (red). Electrodes with significant gamma responses aligned to choices outlined in cyan. Red crosses indicate reference electrodes **C.** Group level one-sample t-test results from a GLM analysis. Significant clusters visible in choice aligned results for reward and the value of the current and counterfactual rule (*top*, *middle* and *bottom* panel respectively, threshold z > 2.33 and cluster corrected at p<0.05). The spectral content associated with reward delivery was primarily observed in the gamma frequency bands, whilst counterfactual rule was associated with bursts of gamma and beta activity. Summary average spectra for reward (dark grey) and counterfactual value (light grey) obtained from the group analysis calculated from 250ms of post choice activity shown on the right (see Materials and Methods for details). **D.** Rule-error aligned, post-choice LFP activity in the beta (blue, 15-30 Hz), low gamma (grey 30-45 Hz), and gamma bands (red, 55-95 Hz). Activity shown relative to either an initial rule error (i.e. error preceded by a minimum of 5 correct trials; left two panels) or a final rule error (i.e. error followed by at least 5 correct trials; right two panels).

Averaging of the activity observed in FPC during shape and colour trials, revealed no significant difference in the LFP activity associated with the two abstract rules (Fig S7). Therefore, we conducted a session-by-session GLM analysis to link the choice aligned spectrogram with time-series encoding behaviourally relevant information. Given previous reports emphasising the importance of transient changes in the LFP activity recorded from the prefrontal cortex [26], this GLM analysis compared fluctuations in LFP activity, on a trial-by-trial basis against the values obtained from the RL models. In line with the hypothesised role of FPC, this analysis included regressors relevant both to exploitative and explorative behaviour namely: reward delivery, the predicted value of either the current and counterfactual rule, as well as the prediction error of both rules (Fig S8, see Materials and Methods for details). Spectrograms displaying the t-stats obtained from the group level analysis revealed significant activity in both the low (30-45 Hz) and high gamma frequency (55-95 Hz) bands, associated with reward delivery. The onset of this activity coincided with animal’s choice and extended for a further 300ms, corresponding to the period during which animals received rewards for correct choices (Contrast: Reward; peak z-stat: 4.92, 93 Hz and 268 ms, Fig 2C). In addition, these analyses revealed bursts of activity, occurring predominantly 400-600 ms after the animals made a choice which were significantly associated with the predicted value of the counterfactual rule. Although the peak of this activity was in the high gamma frequency band (Contrast: Counterfactual Value; peak z-stat post choice: 3.92, 80 Hz and 433 ms), it extended to lower frequencies and included a strong beta frequency component. Analysis of the activity observed in both animals confirmed this observation. Whilst gamma frequency activity (including both low and high gamma activity) was associated with both reward delivery and counterfactual value, beta frequency activity was only associated with counterfactual value (Fig S9). Finally, there was no evidence that choice aligned neuronal activity encoded the predicted value of the current rule (Fig 2C), nor the prediction errors obtained from the RL models which related to either the current or counterfactual rules (Fig S8).

To better understand the relationship between LFP activity recorded from FPC and the animals’ decisions we performed two additional analyses. Firstly, we examined changes in the power of beta, low and high gamma frequency activity relative to rule errors made by both monkeys (Fig 2D). Secondly, we analysed inter-trial activity in FPC to determine whether there was any relationship between persistent, background activity in FPC and counterfactual value (Fig 3).

**Fig 3.**
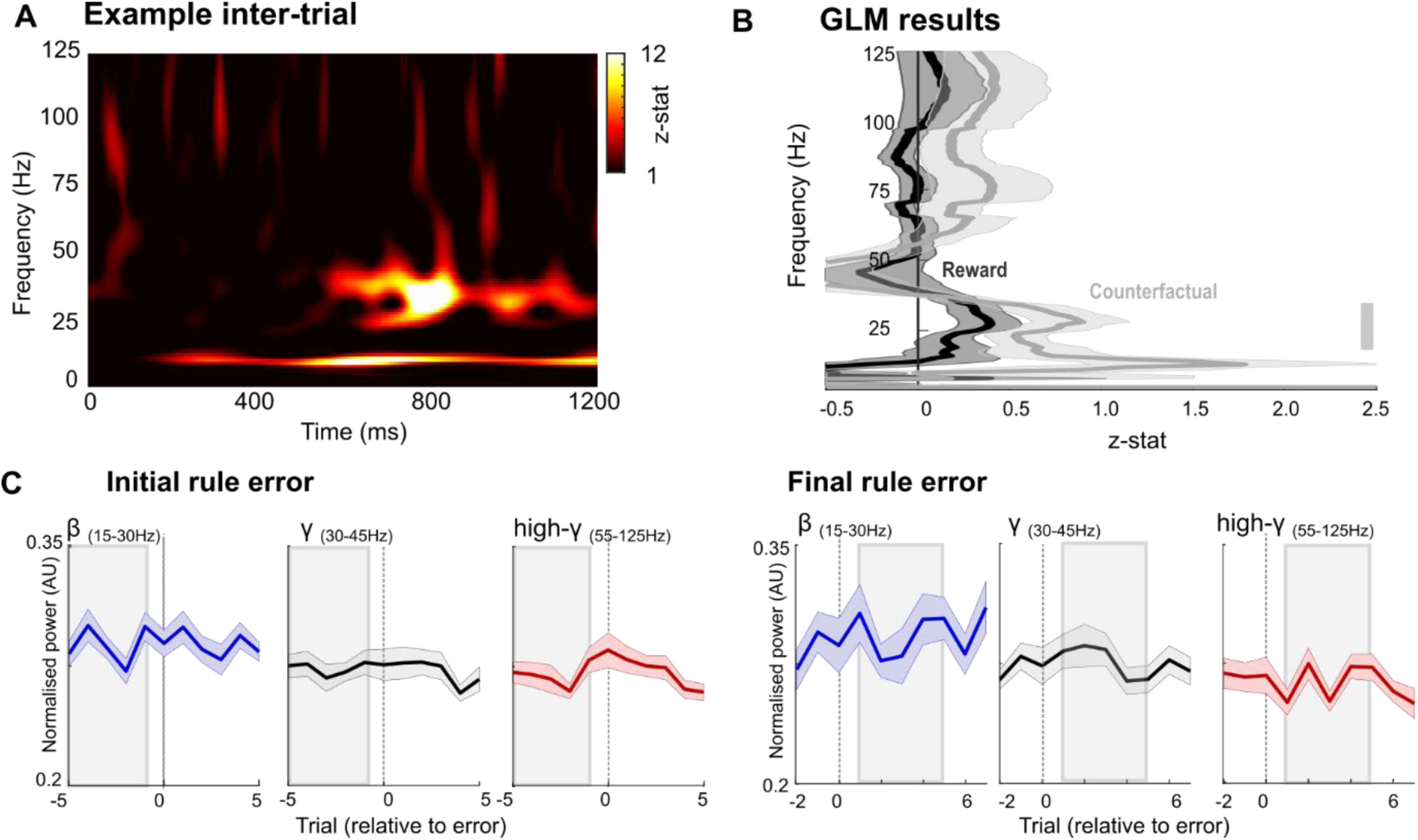
Inter-trial interval field potential activity in FPC. **A.** Example spectrogram from a single period of 1.2 sec of inter-trial interval activity from FPC, taken directly after the end of the trial (See Materials and Methods). LFP activity in FPC dominated by low frequency activity (<50 Hz) **B.** Power spectra showing the results of two contrasts (counterfactual value, grey; reward, black) from a session-by-session GLM analysis (see materials and methods). Significant association shown between counterfactual value and beta frequency activity shown by grey bars (p<0.05, one-sample t-test, cluster corrected at p<0.05). **C.** Rule-error aligned, inter-trial LFP activity in the beta (blue, 15-30 Hz), low gamma (grey, 30-45 Hz) and high-gamma bands (red, 55-95 Hz). Activity shown relative to either an initial rule error (i.e. error preceded by a minimum of 5 correct trials; left two panels) or a final rule error (i.e. error followed by at least 5 correct trials; right two panels).

The power of beta and gamma activity relative to either an initial rule error (i.e. a rule error preceded by at least 5 correct trials) or a final rule error (i.e. a rule error followed by at least 5 correct trials) was highly dissociable (Figure 2D). Beta activity in FPC peaked around the time of the feedback immediately following a rule error (i.e. on trial zero in Figure 2D). Furthermore, the magnitude of feedback-associated beta activity increased across runs of consecutive correct trials (observed both in the run up to an error shown in initial rule error plot, and after a rule-error shown in final rule error plot). In stark contrast, the power of high gamma activity in FPC was lowest in the feedback epoch immediately following a rule error, and during runs of consecutively correct trials high gamma activity remained consistently high. A similar pattern was evident for low gamma activity, which was lowest for rule errors, and stronger for correct trials (although with greater variability than observed for high gamma activity (Fig 2C).

Taken together, these observations, that post-choice beta frequency activity in FPC follows the counterfactual value obtained from RL models (Fig 2C), and that peak power increases over consecutive correct trials (Fig 2D), suggest that beta frequency activity within FPC may coordinate long-term maintenance of value representations between decisions. To evaluate the generality of the timing of this relationship we examined LFP activity recorded from FPC during the corresponding intertrial intervals (i.e. an epoch 1.2 seconds after the end of the trial, see materials and methods, and Fig 3). The LFP recorded from FPC during the inter-trial interval was dominated by activity in lower frequencies (< 50Hz, Fig 3A). Consistent with the above results, linking choice aligned beta activity with counterfactual value, GLM analyses of activity during this intertrial period revealed a relationship between beta frequency activity (15-30 Hz) and counterfactual value (peak z-stat 2.73, 22 Hz Fig 3B). However, analyses of changes in the amplitude of this activity relative to rule errors revealed no direct relationship between either beta, low or high gamma frequency activity and the errors or correct choices made by the animals (initial rule error and final rule error plots; Fig 3C). These findings suggest that whilst the amplitude of offline (intertrial interval) beta activity in FPC does contain information relating to counterfactual value, it is of no direct relevance to the animals’ decisions (correct or otherwise). Importantly, this implies the within-trial-related activities (Fig 2C,D) are not sustained activities maintained across the inter-trial interval, rather they are dynamically reconfigured around the time of decisions.

### High frequency micro-stimulation of FPC impairs adaptation to rule changes and perturbs local field potential activity in FPC

The preceding data demonstrate the co-existence of two task parameters evident in the LFP recorded from FPC. Whilst not completely dissociable, our data suggest gamma activity was associated with reward delivery (and a correct trial outcome), whilst beta frequency activity tracked the predicted value of the counterfactual rule. These correlational observations support the hypothesis outlined earlier that FPC is involved in redistributing resources towards alternative goals (i.e. to counterfactual rules in this case). To directly determine causal links between FPC and decision making we next used electrical micro-stimulation via the implanted electrodes. Microstimulation at high frequencies has previously been observed to perturb both the physiological function of local circuits and emergent cognitive processes [27, 28]. Based on these studies and our aforementioned observations of gamma activity within FPC we chose a “high frequency” stimulation protocol with a peak stimulation frequency of 75 Hz, consisting of 16 pulses, with alternating inter-pulse intervals of 13ms (75Hz) followed by 40ms intervals (Fig 4A & B, and Materials and Methods for more details). Micro-stimulation was delivered on a single trial within each ‘stimulated block’ and was triggered by the reward delivery once animals attained >85% across 10 trials (hence typically occurring ∼8-10 trials prior to the next rule reversal, see Fig 4 and Materials and Methods). Blocks to be stimulated were chosen pseudorandomly and approximately 40% of blocks within stimulation sessions had this single trial stimulated therein, with the rest remaining non-stimulated control blocks. Spectral analysis of the stimulation artefact generated within FPC confirmed that this stimulation caused two pronounced frequencies of activity in the LFP, one at 75Hz and one at 25Hz (Fig 4B).

**Fig 4.**
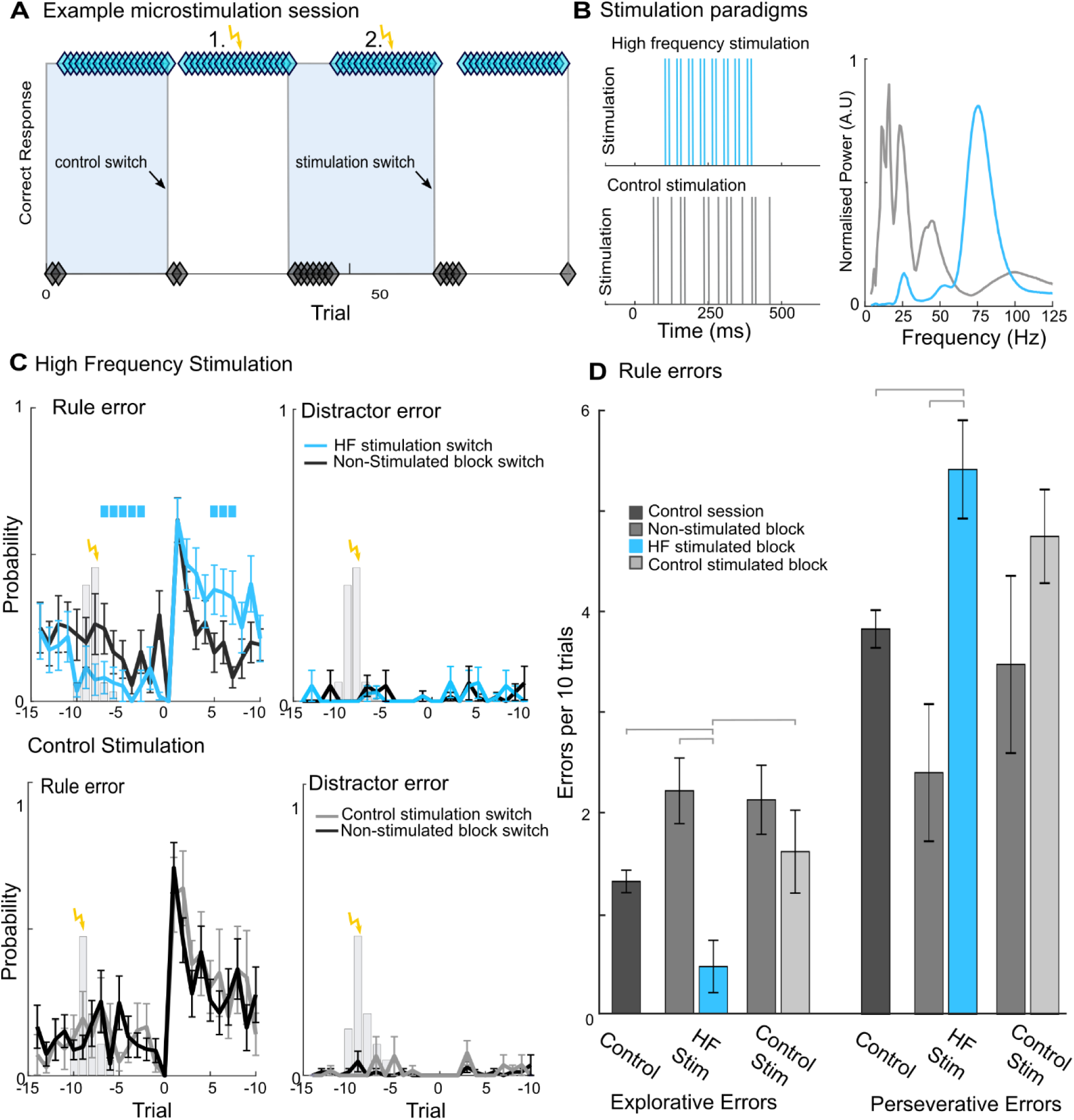
Modulation of monkeys’ behavioural performance by high-frequency electrical micro-stimulation of FPC. **A.** Illustration showing the stimulated trials within an example micro-stimulation and recording session. Micro-stimulation was triggered by the reward pulse, after monkeys had achieved 85% correct over 10 trials (indicated in yellow). Micro-stimulation was delivered to a single trial in a pseudorandom selection of both colour and shape blocks in a session. Post stimulation block changes indicated with black arrows. Diamonds denote correct (cyan) and incorrect trials (black) **B.** Illustration of the two micro-stimulation protocols, high-frequency (*blue)* and control stimulation (*gray, both shown on left*), and the LFP spectral content of both protocols recorded during stimulation from FPC (*right*). See Materials and Methods for details of the stimulation protocols. **C** Plots showing the number of rules errors and distractor errors made by monkeys at the block change following micro-stimulation. High-frequency (*blue*) and control micro-stimulation (*gray, right*) compared to non-stimulated blocks from within the same behavioural session (*black*). All data shown mean of both monkeys ±SE. Significant differences between stimulation and non-stimulation indicated with colour bar, and tested using unpaired t-tests (see Materials and Methods). All p-values corrected using Bonferroni correction. Grey bars denote probability of single-trial stimulation occurring (relative to block change). **D**. Comparison of explorative and perseverative errors made following high-frequency (blue), and control stimulation as well as for non-stimulated block changes within both stimulation sessions and block changes during non-stimulation control sessions. Significant differences determined by Bonferroni corrected post-hoc t-tests.

We hypothesised that this artificial high-frequency stimulation of FPC should affect the region’s influence on cognition, either counterfactual valuation (mediated by bursts of gamma and beta activity) or reward processing (associated with gamma activity alone). Accordingly, these two hypotheses predict contrasting behavioural changes. The first hypothesis supposes disruption to the encoding of the predicted value of the counterfactual rule in FPC. According to this hypothesis the disruptive stimulation would disrupt or suppress the representation of the value of the counterfactual rule, leading in turn to fewer explorative rule errors, and improved behavioural performance in trials that are post-stimulation but prior to the upcoming rule/block change. As anticipatory counterfactual rule-value information is beneficial for rapid switching, this hypothesis also predicts increased perseverative errors at the first post-stimulation rule switch reflecting the need to relearn the information wiped out (or suppressed) by microstimulation. The second hypothesis concerns disrupted reward representations in FPC. According to this hypothesis the stimulation might be expected to impair reward-based rule consolidation before the upcoming rule/block switch thereby increasing uncertainty-triggered exploration (i.e. more explorative rule errors) in the trials running up to the rule/block change. This second hypothesis also predicts impaired performance post-rule/block change because it might be expected to impair reward-based rapid updating of relative rule values. In addition, we utilised a “control stimulation” protocol in which the inter-pulse interval was a pseudo-random integer from 5-100ms. This stimulation protocol was not expected to induce marked disruption to LFP activity within FPC; indeed, analysis of the stimulation artefact associated with this protocol confirmed the absence of high frequency activity (activity extended from 15-45 Hz, Fig 4B).

Our initial analysis of stimulation induced changes in animals’ behaviour examined changes in behaviour following stimulation relative to non-stimulated blocks within the same session. Alignment of the animals’ behaviour with the stimulated trial did not reveal an immediately discernible change in the animals’ behaviour after either stimulation protocol (Fig S10). However, as the animals approached the subsequent rule change marked changes were evident in the probability of monkeys making rule errors following the high-frequency stimulation protocol (Fig 4C).

During the intervening trials post-stimulation but prior to the subsequent rule switch, monkeys made significantly fewer explorative errors following high-frequency stimulation than for non-stimulated blocks (probability of making explorative errors significantly lower from three to seven trials prior to rule switch, p<0.05 cluster corrected, the effect peaked three trials prior to rule switch prior to rule switch, 0.17 vs. 0.05 in no stimulation and high-frequency stimulation respectively, Cohen’s d 1.50, p = 0.004). In addition, there was a significantly higher probability of monkeys making perseverative rule errors in the trials immediately after the first post-stimulation rule switch following high-frequency stimulation than during non-stimulated blocks (probability of making perseverative errors significantly lower than from trials five to seven post rule switch, this effect peaked five trials post rule switch, 0.16 vs. 0.38 in no stimulation and high frequency stimulation respectively, Cohen’s d 1.96, p = 0.04). By contrast high frequency stimulation had no effect on the probability of animals choosing the distractor target either before or after the post-stimulation rule switch (probability of making distractor errors for all trials pre and post rule switch was p>0.05, Fig 4C).

Comparison of the monkeys’ performance following control stimulation protocol (without a high-frequency component, Fig. 4B) with non-stimulated trials from the same session did not reveal any significant changes in the monkeys’ ability to perform the WCST. There was no significant difference between the probability of monkeys making rule errors with control stimulation vs. no stimulation either prior to (probability of making a perseverative error peaked four trials before block change 0.15 in non-stimulation blocks vs. 0.10 in control stimulation blocks, Cohen’s d, 0.58, p=0.33) or following the block change (probability of making a perseverative error peaked five trials after the block change 0.20 in non-stimulation blocks vs. 0.26 in control stimulation, Cohen’s d, 0.59, p=0.30). Consistent with the previous results, control stimulation did not change the probability of animals making distractor errors either before or after the subsequent rules switch (Fig 4D).

Further analysis utilising repeated measures ANOVAs to compare the number of explorative and perseverative errors made by both animals across behavioural sessions confirmed these trends (two within subject factors: Stimulation protocol, 3 levels (HF stimulation, control stimulation and no-stimulation) and BlockType, 2 levels (stimulated and non-stimulated), see Materials and Methods, Fig 4D). There was a significant main effect of stimulation protocol for both explorative errors and perseverative errors (F_(2,26)_ = 15.62, p = 8.18 ×10^−7^ and F_(2,26)_ = 14.17, p = 6.89 ×10^−05^ errors respectively). Furthermore, there was a significant interaction between stimulation protocol and block type for both explorative and perseverative errors (F_(4,52)_ = 21.18, p=2.079 ×10^−10^ and F_(4,52)_ = 37.403, p = 1.014 ×10^−14^ respectively). Posthoc t-tests confirmed that the number of explorative errors made by animals was significantly lower following high frequency stimulation than after control stimulation (p = 0.0052, mean explorative errors 0.47 ±0.26 vs. 1.62 ±0.41 per session for high frequency and control stimulation respectively) or when compared to non-stimulation sessions (p = 0.024, mean explorative errors 1.32 ±0.11 per session for non-stimulated sessions).

By contrast, perseverative errors made after post-stimulation block changes by both animals were significantly higher after high-frequency stimulation than at block changes during non-stimulated sessions (p = 0.035, mean perseverative errors 5.42 ±0.50 vs. 2.40 ±0.68 for high-frequency stimulation and non-stimulated session respectively). Although the animals made more perseverative errors following high frequency than following control stimulation the difference was not significant (p = 0.24, mean perseverative errors 4.75 ±0.47 following control stimulation). Finally, there was no difference between the number of explorative or perseverative errors made during non-stimulated blocks from either stimulation protocol or control non-stimulated session.

Taken together, the pattern of errors shown in these results (decreased explorative rule errors following stimulation up to rule-change, coupled with increased perseverative errors after the rule-change, and no evidence of increased distractor errors) is consistent with our first hypothesis that high-frequency stimulation disrupts FPC-mediated behaviour based on its encoding of the predicted value of the counterfactual rule. Finally, to determine if our electrical micro-stimulation protocols caused an enduring affect upon the LFP activity in FPC concomitant with the behavioural change (associated with both hypotheses), we compared changes in LFP activity recorded from FPC after both of the electrical microstimulation protocols (Fig 5). This analysis compared 400ms of choice aligned activity for 10 trials before and 10 trials after the first post-stimulation block change (see Fig 5A & B and Materials and Methods) with corresponding activity recorded before and after non-stimulated block changes from within the same session. A repeated measure ANOVA with one between subjects factor: Monkey (two levels, M1 and M2) and one within subjects factor: Stimulation (two levels, stimulation and non-stimulation) revealed significant differences between LFP activity recorded following high-frequency micro-stimulation and non-stimulated activity (Fig 5C). This analysis revealed two clusters which were significantly different between high-frequency stimulated and non-stimulated trials. Firstly, following high-frequency micro-stimulation activity in the high gamma activity (55-95 Hz, peak z-stat 2.81, 90 Hz, three trials pre-block change) was significantly lower than in non-stimulated blocks prior to the subsequent rule-switch and associated block change (trial 0). In addition, a second cluster of decreased activity was evident post rule-switch. This cluster was evident over the first eight trials after the change in rule and was predominately in the low gamma frequency (z-stat 3.08, 24 Hz, one trial post rule switch) and beta frequency band (peak, z-stat 3.74, 44 Hz, eight trials post rule switch), although it extended into the high gamma frequency bands immediately after the rule switch (Fig 5C). By contrast a comparable analysis between control stimulation, and non-stimulation trials did not reveal any significant differences in field potential activity recorded from FPC, either before or after the block change (Fig S11).

**Fig 5.**
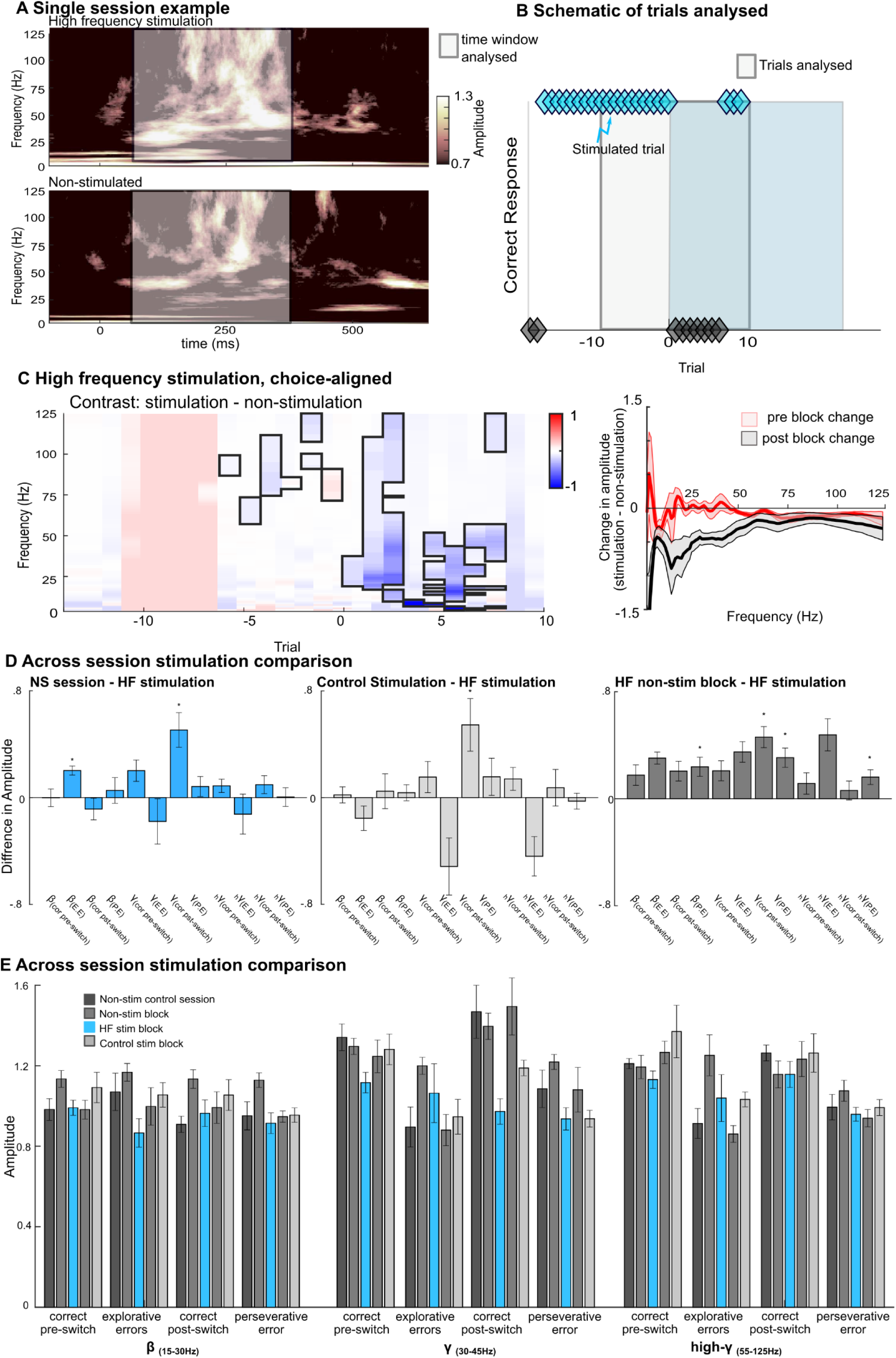
Disruption of choice-aligned LFP activity in FPC following high-frequency electrical micro-stimulation. **A.** Example spectrograms showing choice-aligned activity recorded from FPC following high-frequency stimulation (*upper*) or in the absence of stimulation (*lower*). Spectrogram calculated by averaging over the fifth to eighth trials after stimulation. Non-stimulation spectrogram calculated from the average spectrograms of trials from the last four trials of four non-stimulated blocks (i.e. same relative block epoch). White box indicates the 400 ms period used for further analysis. **B.** Illustration of the trials included in the analysis of changes in LFP activity recorded from FPC caused by electrical micro-stimulation. In stimulation blocks the 10 trials before and 10 trials after the post-stimulation block change were included (white box). In non-stimulation blocks a corresponding window of trials was selected. **C.** Trial by-trial **s**pectral analysis showing the difference between the field potential in FPC following high-frequency stimulation and non-stimulated blocks within the same session **D & E**. Cross session comparison showing **D.** the difference in amplitude between non-stimulated control sessions and high-frequency stimulation (*left*), control stimulation sessions and high-frequency stimulation (*middle*) and between non-stimulated blocks and high-frequency stimulated blocks (*right*) split for beta (β) (15-30 Hz), gamma (γ) (30-45Hz) and high gamma (_h_γ) (55-95 Hz) bands and for explorative (E.E) and perseverative (P.E) errors as well as correct trials pre block switch (cor pre-switch) and correct trials post block switch (cor pst-switch). **E.** LFP amplitude for non-stimulation control sessions (dark grey), stimulated blocks from high-frequency (blue) and control stimulation (light grey), as well as non-stimulated blocks from both stimulated sessions (grey). Amplitudes are shown using the same split as used in D.

To confirm these changes in activity within FPC we compared LFP recorded from FPC following both stimulation protocols within activity recorded from non-stimulated control sessions (Fig 5 D&E). Consistent with results presented in Fig 2 these data (Fig 5E) revealed strong gamma and high gamma responses and low beta frequency activity when the animals chose correctly; by contrast errors (both explorative and perseverative errors) were typified by both lower gamma and lower high gamma responses but increased beta frequency activity. Comparison of the difference in LFP amplitude between non-stimulation sessions and high-frequency stimulation sessions (Fig 5D left panel) revealed that the stimulation decreased the amplitude of activity in FPC for several trial types. Following high-frequency stimulation, but prior to the post-stimulation block change beta activity recorded from FPC for explorative errors was significantly lower than corresponding trials recorded from non-stimulation sessions (difference between non-stimulation and high-frequency stimulation explorative errors 0.2 ±0.03, p=0.0042 one-sided t-test, Holm-Bonferroni correction for multiple comparisons). In addition, low gamma amplitude for correct trials made post-stimulation but pre-block change was lower than for non-stimulation sessions, although not after correction for multiple comparisons (difference between non-stimulation and high-frequency stimulation correct trials pre-block change 0.2 ±0.08, p=0.045 one-sided t-test, uncorrected). Furthermore, gamma activity recorded during correct trials following a block change was significantly lower after high-frequency stimulation than for corresponding trials from non-stimulation sessions (difference between non-stimulation and high-frequency stimulation correct trials post-block change 0.51 ±0.13, p=0.004, Holm-Bonferroni correction for multiple comparisons). The amplitude of gamma activity recorded during correct trials post-block change was also significantly lower following high-frequency stimulation than following control stimulation (difference between control stimulation and high-frequency stimulation for correct trials post-block change 0.55 ±0.2, p=0.0045 one-sided t-test, Holm-Bonferroni correction for multiple comparison, Fig. 5D middle panel).

Comparison of LFP activity following high-frequency stimulation and non-stimulated blocks from within the same sessions (Fig 5D right panel) revealed a comparable decrease in gamma for correct post-stimulation trials (difference between non-stimulated blocks and stimulated blocks from high-frequency stimulations sessions for correct pre-block change trials 0.43 ±0.7, p=0.0001 one-sided t-test, Holm-Bonferroni correction for multiple comparison), as well as a drop in the amplitude of LFP amplitude across all bands for perseverative errors (difference between non-stimulated blocks and stimulated blocks from high-frequency stimulations sessions for perseverative errors 0.24 ±0.07 p=0.045, 0.25 ±0.07 p=0.002, 0.13 ±0.03 p=0.001 for beta, gamma and high-gamma respectively. All one-sided t-test Holm-Bonferroni correction for multiple comparisons).

## Discussion

It has been proposed that FPC’s contribution to decision-making in primates is to track the relative value of alternative options, modulate exploratory versus exploitative tendencies, and so manage competing goals [2]. Little is known about the neural mechanisms that underlie such cognitive processes. Here using a WCST analogue task we present evidence of an association between rhythmic activity in FPC and two key variables needed to guide exploratory rule-guided decision-making, namely counterfactual rule value and reward delivery. Furthermore, we demonstrate causally that temporary disruption of FPC (using targeted electrical microstimulation) affects both the explorative decision-making behaviour of the animals and the concomitant LFP activity in FPC. Specifically, brief FPC stimulation initially made animals better at the WCST task (due to reduced explorative errors) in the run up to a rule-change, before subsequently impairing task performance after the rule changed (due to increased perseverative errors). Importantly, the behavioural effect of stimulating during a single trial could still be observed up to around 20 trials later, and was accompanied by persistent changes in the amplitude of low gamma and beta activity in FPC. Taken together these results not only provide confirmation that FPC is essential for guiding exploratory decision making but yield mechanistic insights into how neuronal activity within FPC can support rule-guided decision-making behaviour. Both of these stimulation-induced behavioural changes were consistent with the proposed role of FPC in contributing to guiding exploratory decision-making. Firstly, the decreased explorative behaviour prior to the rule-change supports correlative evidence from human neuroimaging which links activity in FPC with switching behavioural choices [6, 29] and exploratory decision-making [30, 31]. Secondly, the increased perseverative error-rate after the rule/block change is consistent with patients’ perseverance with previously relevant abstract rules, as well as a slow adaption to new information following damage to or disruption of human FPC [12, 32].

In addition, we present evidence that at least two forms of dissociable information can be observed in the LFP recorded from FPC. Gamma activity (extending across both the low and high gamma band) was directly associated with external information about successful trial outcome, whilst beta activity tracked the subjective (internally generated) value associated with choosing the counterfactual rule. A similar dissociation, between incoming sensory information and internally generated working memory, has been reported in the oscillatory activity recorded from lateral prefrontal cortex (LPFC); in macaque LPFC, gamma activity is pronounced during encoding phases of working memory tasks and is associated with incoming sensory information [33]. Comparable recordings from human LPFC have also reported gamma activity when remembering lists, where such activity scales with the information load of the items to be retained [34, 35]. By contrast, beta activity in LPFC peaks in the absence of sensory information (e.g., during delay periods of working memory) [26, 33, 36] and has been proposed to mediate top-down, internal signals [37, 38], for example strong beta frequency coherence directed from frontal to parietal lobes is a feature of directed spatial attention [39].

Here both animals succeeded in the WCST by periodically exploring throughout blocks. This strategy likely developed through the animals being sufficiently well-trained to possess both a firm concept of the two abstract rules themselves (demonstrated by both animals rarely making distractor errors), and an understanding that the current counterfactual rule will become valuable as more reward is gained from the current rule (see Supplemental Material for further discussion of the animal’s behavioural strategy). Within this framework, the co-existence of both internally generated predictions and sensory information within FPC is particularly notable. Here after a correct choice we observed a consistent high-gamma frequency, and strong if noisy low-gamma response, together with a variable beta frequency component which increased in amplitude across runs of successive correct trials and peaked after rule errors. Interestingly, high-frequency microstimulation did not selectively disrupt either the gamma or beta LFP component as we hypothesised. Rather following microstimulation we observed decreased low-gamma (following correct trials) and beta activity (following explorative errors), raising the possibility that activity in both bands is linked during decision making.

Previous studies have suggested that one feature of neuronal activity in macaque FPC is the evaluation of internally generated decisions [15, 16] during task feedback. The combined nature of the gamma-beta response observed here could reflect the mechanism by which information regarding task success, relayed within the incoming gamma activity, is used to update the persistent counterfactual value in turn organised by beta frequency activity in FPC. Hence a potential interpretation is that micro-stimulation impaired the updating of the counterfactual value within FPC leading to a degraded (or unreadable) counterfactual rule value, resulting in the animals performing better towards the end of blocks as current rule not competing with counterfactual rule. By extension, this diminished counterfactual representation delayed adaptation to the subsequent rule change as the animals required longer to re-acquire and establish the value of the now relevant (previously counterfactual) rule. Importantly, whilst this choice-aligned activity in FPC was affected by microstimulation across several subsequent trials, persistent inter-trial activity was unaffected, suggesting that whilst the value associated with the alternative goal or option is updated within FPC following a decision, this information may then either be communicated immediately to other cortical areas needed for decision making (and only later reconstituted in FPC) or else stored in FPC an activity silent manner analogous to that proposed for working memory [40].

The transience we observed here is consistent with the majority of previous neuronal recordings from FPC which have reported task relevant information (needed to guide future decisions) in the firing of FPC neurons only during the delay/reward feedback [15, 16]. By contrast, recent findings from an object-in-place task found neuronal firing in FPC peaked during the initial presentation of a novel scene, with limited firing for subsequent presentations or during the delay or choice epochs [19]. One parsimonious explanation for these findings is that FPC is only engaged when the relative value of counterfactuals choices is uncertain or in flux. Hence, in associative learning tasks (including the object-in-place task) the relative value of correct versus incorrect (i.e. counterfactual) targets is unchanged after the initial fast-learning phase (during which representations of relative values of novel stimuli are being discovered and accordingly susceptible to significant updates). With subsequent repeated presentations the task becomes an exercise in strengthening object-place associations, without the need to monitor counterfactuals or engage FPC, therefore the presence or complete absence of FPC becomes irrelevant for task performance [8].

In humans, activity relating to the value of alternative choices appears to be localised to the lateral subdivision of FPC [2, 5, 29], distinct from more general feedback related activity in the medial sub-division. Here we recorded from the most anterior-lateral region of macaque FPC and found activity related to both the value of the counterfactual rule as well post-choice feedback, suggesting that when compared to humans, these functions may not be as regionally segregated in the macaque FPC. However, understanding of the functional homologies between human and macaque frontal pole remains nascent, and the degree of overlap in the behaviour supported by FPC in both species remains to be determined. For example, the lateral subdivision of human FPC is functionally connected to a distinct network of regions in a pattern which may be unique to humans [41]. Furthermore, this lateral FPC region is reliably activated in humans during complex behaviour such as cognitive branching [2, 5]. Cognitive branching requires a more structured consideration of counterfactual task rules (i.e. retaining knowledge of which counterfactual rule to return to) and it is not clear whether macaques possess the cognitive ability to perform this highest level of directed exploration (see supplementary material for further discussion of functional homologies between human and macaque FPC).

Dysfunction of FPC is common in patients with obsessive compulsive disorder (OCD), where under-recruitment of this lateral subdivision into the cortical networks supporting decision-making has been associated with deficits in set-shifting [42], task load [43], and emotional decision-making [44]. By contrast, FPC hyperactivity is observed in ADHD [45]. Indeed, significant developmental changes in FPC are associated with disturbed behaviour in both ADHD [46] and Schizophrenia [47]. The nature of neuronal activity within human lateral FPC remains to be determined, however our results suggest that coordinated neuronal activity at beta frequency may be crucial to normal functioning of FPC. Future research characterising the dynamics between FPC and the wider network of cortical areas with which it is connected [48, 49], will be essential to understanding how maladaptive cortical networks involving FPC can influence decision making as well as targeting interventions to ameliorate sub-optimal decision-making.

## Materials and Methods

### Animals

We recorded the LFP from, and performed electrical micro-stimulation of, fronto-polar cortex (FPC) in two young male macaque monkeys (*Macaca mulatta*) whilst they performed the WCST analogue [22, 50]. At the time of recording monkeys were aged 8 and 9 years, and weighed between 10 to 12 kg. Both monkeys were socially housed in enriched environments with a 12hr light/dark cycle and had *ad libitum* water access. All animal surgery, anaesthesia, and experimental proce-dures were carried out in accordance with the guidelines of the UK Animals (Scientific Procedures) Act of 1986, licensed by the UK Home Office, and approved by Oxford’s Committee on Animal Care and Ethical Review.

### Apparatus

The WCST analogue was programmed using Turbo Pascal (Borland), run under DOS on a desktop PC. The animal was sitting in a primate chair (Rogue Research Inc.) in front of the touch screen with the head-fixated and performed the task in a magnetic-shielded box. A window in the front of the chair provided its access to the touch-screen and a touch-sensitive knob which we refer to as a ‘key-touch’ device that the animal had to steadily hold at various times in each trial and which was positioned low down in front of the touch screen. A peristaltic pump device located on top of the box fed trial-by-trial smoothie reward through a tube and to a spout positioned in the vicinity of the animal’s mouth. Below the screen was also an automated lunch-box which contained the majority of the animal’s daily meal (wet mash and fruits and nuts etc.) and which only opened immediately at the end of the task thereby providing an additional and main motivation to the NHP to complete the session sooner rather than later (as can be achieved by good and focussed task performance).

### Behavioural Task

Prior to training on the WCST analogue both monkeys underwent preliminary training and shaping, consistent with procedures described previously [22, 50]. Both monkeys were then trained to per-form the WCST macaque analogue as previously described in detail (see [7, 22] for full task-train-ing stage-by-stage details). The WCST task used in this study used 36 different stimuli comprising six colours (red, yellow, green, cyan, blue, and magenta) and six shapes (square, triangle, circle, hexagon, cross, and ellipse). The sample in each trial was selected at random (without replacement until the entire set had been used) from the 36 stimuli. In each trial the test items were also selected from the same set of 36 stimuli, and at random (with the restrictions imposed by the necessity to generate either a low- or high-conflict trial). The locations of the three test items (i.e., to the left/right/bottom of the sample) were also chosen at random. Together these stimulus selection pro-cedures ensured that to succeed at the task the animals could not learn about rewarded objects, ob-ject-features, or object-location; rather they had to learn abstract matching rules (see below).

Briefly, on each trial of the WCST analogue, monkeys were presented with one central sample stimulus, and following a delay three peripheral choice stimuli. One of the choice stimuli matched only according to colour of the sample, and another matched only according to shape of the sample, while the third-choice stimulus shared no features. On a block-by-block basis the correct choice was determined by either colour or shape. The monkeys received no cue about the currently reinforced rule and so had to determine the correct rule based on feedback from preceding trials. The rule switched periodically but only after monkeys achieved performance >= 85% correct in 20 consecu-tive trials. Prior to electrode implantation surgery both monkeys received extensive training on the WCST analogue, with additional training post-implantation. For recording only sessions of this study (collected at the beginning of the study) the animals completed an average of 5 blocks, (i.e. rule reversals) per recording session (6.25 for M1 and 3.5 for M2).

### Neuronal recording and electrical micro-stimulation

In addition to a titanium mount for head restraint (Gray Matter Research, Bozeman, MT) each mon-key was implanted with a total of four or five Utah arrays (Blackrock Microsystems, Salt Lake City, UT) with platinum iridium tipped electrodes. These arrays were connected to two 128 channel ped-estals (256 channels in total) and arrays were implanted into FPC and orbitofrontal cortex (OFC), and dorsal and ventral lateral prefrontal cortex (dlPFC and vlPFC); M1 received an array implanted in ACC. The location of the arrays implanted into FPC were confirmed by examination of the brain post-mortem and are shown in Figure S4. At the time of this microsimulation study (16 months af-ter implantation in M1 and 11 months after implantation in M2) neuronal activity was only consist-ently observed in both animals in the FPC arrays (a 32 channel Utah array in M1 and a 64 channel Utah array in M2). Neuronal data was collected using a 256 channel Cerebus System and concur-rent electrical microstimulation was carried out using a CereStim 96 system (all Blackrock Mi-crosystems Salt Lake City, UT).

The raw wideband LFP was sampled at 30 kHz and saved offline for further preprocessing. These data were then downsampled to 1kHz and bandpass filtered between 4–150 Hz. Mains noise was removed by notch filtering using an equiripple FIR filter with an attenuation of 30dB between fre-quencies from 48-52Hz, surrounded by a ±1 Hz transition band. Any residual harmonics were removed using comparable filters at 100, and 150 Hz. After removal of mains noise data from all electrodes were visual inspected to ensure signal quality. Electrodes with suspect signal quality (6 electrodes from M1 and 34 electrodes from M2) were not included in further analysis. Data pre-sented in this study were collected from 18 recording only sessions (monkey 1: 8 sessions, monkey 2: 10 sessions) and 35 recording and stimulation sessions (monkey 1: 18 sessions, monkey 2: 17 sessions).

Electrical microstimulation consisted of trains of fourteen 100 μA, 200 μs biphasic pulses (cathodal first). Two different stimulation protocols were delivered to examine FPC function. During the high-frequency protocol, the inter-pulse-interval alternated between 13 ms and 40 ms, this protocol induced a pronounced 75 Hz response in the LFP recorded from FPC (Fig 3). In the control stimula-tion protocol, the inter-pulse-interval was a pseudo-random integer from 5-100ms. This stimulation protocol caused a strong low, but not high frequency response, in the LFP recorded from FPC (Fig 3). On each pulse 3 electrodes were stimulated, the exact combination of electrodes varied on each pulse between 6 electrodes pre-selected with strong gamma frequency activity aligned to reward de-livery. In micro-stimulation sessions, the stimulation was delivered approximately half the blocks in the session, the blocks stimulated were selected pseudo randomly. Microstimulation was never de-livered on the first block of the session. Overall microstimulation was delivered on approximately 40% of blocks in each session (stimulation delivered on average on 3.1 and 1.3 blocks out of 8.1 and 3.3 total blocks per session for M1 and M2 respectively). Stimulation was delivered on a cor-rect trial once monkeys had achieved 85% correct over 10 trials and was triggered by the reward delivery.

### Data analysis

All data analysis and pre-processing was carried out using freely available toolboxes and bespoke scripts written for MATLAB 2018B (The MathWorks Inc., Natick, USA). Data was imported to MATLAB using the NPMK toolbox (4.4.0.0), and spectral analysis of LFP was carried out using the FieldTrip toolbox [51].

### Behavioural modelling and analysis

To aid our initial analysis of the animal’s behaviour, we classified the monkeys’ incorrect responses into several error categories. When animals chose the stimulus corresponding to the incorrect/alter-native rule these trials were deemed “rule errors”. We further use the term perseverative and explor-ative error to refer to rule errors made during the start of the block (when animals perseverated with the previously correct rule) and at the end of the block (when animals continued to explore the in-correct rule) respectively. For the purpose of the RT analysis in Figure S2 and the comparison of model predicted explorative errors these errors were deemed to be rule errors made in the final 7 trials of the block.

To determine the cause of the explorative errors made by monkeys whilst performing the WCST analogue the value attached to both matching rules (colour and shape) was modelled with four vari-ants of a reinforcement learning (RL) model [24, 25]. The first two models (RL _1 chosen_ and RL _2 chosen+ unchosen)_ were included to test the hypothesis that the monkeys tracked both the current and counterfac-tual rule. In addition, two models were fit to test alternative hypotheses for the generation of explor-ative errors. Firstly, that explorative errors occurred because the animals had difficulty recalling the correct rule towards the end of blocks (RL _3 random noise_). Secondly that these errors simply reflect the animals becoming distracted on individual trials (RL _4 point disruption_).

In the first model (Fig 1 D, RL _1 chosen_) only the value of the rule corresponding to the chosen target (and not the alternative) was updated. In this model, if on trial *n* the monkey chose shape then the value *vC* of the colour rule was unchanged whilst the value *vS* of the shape rule for trial *n + 1* was given by.

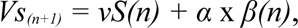

Where the constant *α* was the learning rate, and *β(n)*, the prediction error, was given by.

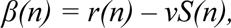

Where *r(n)* was the reward obtained for trial *n.* The same procedure was followed when monkeys chose colour to calculate *vC*, the value associated with colour. Both values were unchanged on trials in which the animals chose the other rule. For example, on a trial in which the monkey correctly chose to match to shape, only *vS* will be increased, with *vC* was unchanged as it wasn’t chosen. On the rare occasions that the animals chose the distractor (the target which matched neither shape nor colour), neither value was updated.

In the second model (Fig 1 D, RL _2 chosen+ unchosen_) the value of both the chosen and unchosen rules were updated according to the above equations on every trial regardless of which rule was chosen. That is, on an example trial in which the monkey correctly chose to match to shape, both *vS* and *vC* will be updated, with *vS* and *vC* increasing to reflect the outcome of the trial. Note that the equations used to calculate the chosen (here *vS)* and unchosen (*vC)* were independent, and that two separate learning rates, *α,* were utilised, with the learning rate assigned to the counterfactual rule limited to a range of 0.005 −0.5.

For the third model, (Fig S1 A, RL_3 random noise_) only the value of the chosen rule was updated on each trial, the value of the unchosen rule was left unchanged (as above for RL _1 chosen_). To represent the failure of the animals to assign value to the correct rule as blocks progressed, the updated value (*vS* or *vC*) was modified by a randomly generated decimal, which increased over the length of the block.

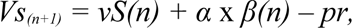

Where *pr* was a randomly generated value from 0-1, which increased with block length, normalised so that 1 was equal to the length of the longest block in each session (see Fig S1 for an example session of the random value).

The final RL model (RL _4 Point distraction_) was designed to capture the hypothesis that explorative errors arise from the monkeys incorrectly allocating value to the wrong rule for a single trial within the task (e.g., becoming distracted for a single trial). Similar to RL _1 chosen_ this model updated only the value of the chosen rule on each trial. However, the value of the chosen rule was switched with the unchosen value for single trials selected pseudo-randomly. Point Distraction trials were selected when an accumulation function, which increased from 0-1 during the block crossed a stochastic threshold, which varied between .75 and .95 (an example of the threshold and accumulation function shown in Fig S1 B).

For all four models the probability of choosing either rule was then estimated from the two values (*vS* and *vC*) using a softmax function. For example, the probability of choosing shape (*pS*) on trial *n* was given by.

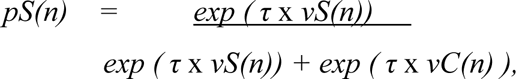

Where the constant τ, inverse temperature determines the randomness of the decision.

### Model optimisation and comparison

The optimum values for the constants *α* and τ were obtained by finding the combination of parameters which minimised the negative log likelihood of the predicted choices for each model.

To determine the model which best fit the monkeys’ actual choices the Akaike Information Crite-rium (AIC) was calculated from the minimised negative log likelihood obtained for the four models for session (for reference the AIC values for all four models, and AIC_null_ for all sessions are shown in Fig S3). Goodness of fit was calculated using Mcfaddens Pseudo-R^2^ [52, 53] a metric summaris-ing goodness of fit, derived from the AIC for which models with pseudo-R^2^ values between 0.2-0.4 are considered to fit the observed data well [53].

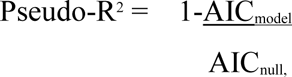

Where AIC_null_ was the AIC calculated from a null hypothesis that monkeys chose randomly between the three available choices (match shape, match choice and distractor) on each trial [52, 53].

A repeated measure ANOVA was calculated to assess differences in model fit between the four models. This ANOVA had one between level factor, Monkey (two levels; m1 and m2), and one within level factor, Model (four levels; RL model _1-4_). Post-hoc t-tests, corrected for multiple com-parisons using Bonferroni correction, were used to assess differences in the model fit overall and for each monkey separately (data split by monkey are shown in Fig S3 B).

Finally, session by session GLMs were used to confirm that improvements in fit in the RL model were due to better prediction of the explorative errors made during the final trials of a block. For each session a single time series was created, corresponding to the observed probability of explora-tive errors being made during the 7 trials preceding a block change. A GLM use then used to com-pare this time-series with the fit of predictor time series derived from the corresponding sections of predicted perseverative errors from all four RL-models (RL _1-4_). The GLM results from individual sessions were combined using a further, second-level, repeated measure ANOVA with had one be-tween level factor, Monkey (two levels; m1 and m2), and one within subject factor, Model (four levels; RL model _1-4_).

Within session differences in the behavioural performance of M1 and M2 following electrical mi-crostimulation were assessed by examining the perseverative errors made by both monkeys, aligned to the subsequent block change. Unpaired t-tests were used to assess the probability of monkeys making perseverative errors in stimulated and non-stimulated trials for the 7 trials before and after a block change. Corrections for multiple comparisons were made using Holm-Bonferroni.

Between session differences in behavioural performance were examined across non-stimulation session, high-frequency stimulation sessions and control stimulation sessions. Data from stimulation sessions were split into stimulated block changes and non-stimulated block changes respectively giving five conditions (non-stimulation sessions, high frequency stimulation sessions stimulated block changes, high-frequency stimulation sessions non-stimulated block changes, control stimulation stimulated block changes, control stimulation non-stimulated block changes) The number of explorative errors (rule errors made over the last 10 trials of a block) and perseverative errors (rule errors made over the first 10 trials of a block) were averaged per session and differences between conditions assessed using two repeated measure ANOVA’s, for explorative and perseverative errors respectively. These ANOVA’s had one between subjects factor, Monkey (two levels; m1 and m2) and two within subject factor, Stimulation type (two levels; stimulated and non-stimulated) and stimulation protocol (three levels; control, high-frequency stimulation and control stimulation). Post-hoc t-tests were carried out using Holm-Bonferroni correction for multiple comparisons.

### Analysis of the local field potential

To link the LFP activity recorded from FPC with behavioural cues we conducted spectral analysis of the activity recorded from FPC. The first step in this analysis was to examine cue aligned spectrograms for every electrode in the arrays implanted in FPC. For each electrode, we selected 1000 ms (−400 ms to +600 ms) segments of LFP, aligned to one of three behavioural cues (sample onset, target onset and choice).

Spectral analysis was carried out on a trial-by-trial basis to preserve induced neuronal activity (i.e. activity occurring with a variable latency after the relevant behavioural cue). Spectrograms were calculated for every trial from the aligned LFP segments using Morlet wavelets with frequencies ranging from 4 Hz to 124 Hz in 1 Hz steps. The full-width at half-maximum (FWHM) ranged from 656 to 123 ms with increasing wavelet peak frequency. In the frequency domain this corresponded to a FWHM from 1.5 Hz to 45.5 Hz (See Fig S12, [54]). Spectral decomposition was carried out using the FieldTrip toolbox (ft_specest_wavelet function,[51]). Induced single-trials spectrograms were then normalised to the mean spectral content of 800 ms of intertrial interval activity prior to the start of each trial (averaged both across all trials, and across the 800ms of activity) before further analysis.

After the initial spectral decomposition, a secondary analysis was conducted to identify electrodes with increased activity associated with either of the three behavioural cues. In this analysis responsive electrodes were identified using paired t-tests to compare the mean activity between 20 and 100 Hz, −400 to −200 ms pre-cue with activity in the corresponding frequency bands 200 to 400ms after the cue.

A session-by-session univariate GLM was used to link reward aligned spectral activity to regressors representing physiologically relevant variables on a trial by trial basis. This GLM included a regres-sor denoting whether monkeys received a reward on each trial as well as four regressors derived from the RL model _2 current + unchosen_ described above. Two of these regressors denoted the estimated value of the current correct rule and the alternative rule (i.e the value of choosing a shape target whilst in a colour block). To improve signal to noise these two regressors for rule value were con-volved with a Gaussian function with a width of 4 trials. Finally, two regressors were included which corresponded to the trial-by-trial difference in the two estimated values (delta of the current and other rules). These time series were rectified to remove negative values and convolved as above. The GLM for each time frequency point was given by:

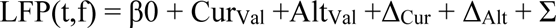

Where LFP(t,f) is a variable containing the amplitude of the rewrad aligned spectrogram for a given time,frequency point across all trials in session. Cur_Val_ and Alt_Val_ are regressors for the trial by trial estimates of the value of the current and alternative rule as described above and Δ_Cur_ + Δ_Alt_ regres-sors corresponding to the un-signed change in the value of each rule as descrcibed above.

This GLM was applied to all time, frequency points in the reward aligned spectrograms for each electrode separately, before averaging across electrodes to yield five parameter estimate (PE) spec-trograms for each session. A one-sample t-test was applied to each time-frequency bin of each PE of the five PE conditions to test the hypothesis that activity in each bin was associated with each re-gressor. The resulting five statistical summary spectrograms (one for each regressor) were then thresholded at z = 2.33 and cluster corrected at p<0.05.

LFP activity during the intertrial interval (absent of task relevant cues or external events) was ana-lysed using a comparable pipeline. For all electrodes included in the choice aligned analysis (se-lected above) a 1.2 sec data epoch a selected directly following the end of each trial. Wavelet de-composition was carried out as described above (including normalisation to the mean spectra of the intertrial-interval). Session-by-session GLM analysis, using the same behavioural regressors, was carried out on the resulting normalised spectrograms. However, to account the highly variable na-ture of intertrial activity, without external events to align activity across trials. This intertrial GLM analysis was only carried out in the frequency domain. Values of the time by frequency spectrograms less than 1 (i.e., low than the session mean intertrial values) were removed from the before averaging across all 1.2 seconds of data in the temporal domain. The GLM was then applied to each frequency bin of the resulting mean power spectrum. As above statistical significance was determined using one-sample t-tests, comparing the parameter estimates from all sessions for a fre-quency bin against a null hypothesis that there was no relationship with each regressor. The result-ing five statistical summary power spectra (one for each regressor) were then thresholded at z = 1.66 and cluster corrected at p<0.05 (again with 1000 random permutations).

### Analysis of changes in LFP activity following electrical micro-stimulation

To determine whether electrical micro-stimulation had an impact on a particular frequency band of field potential activity recorded from FPC, we examined choice aligned LFP activity, before and after the first post-stimulation block change and compared this to corresponding activity recorded from block changes which were not preceded by micro-stimulation (Fig 4A & B). We calculated changes in the LFP caused by electrical micro-stimulation by averaging 400ms of choice aligned spectrograms from 20 trials aligned to either the first post-stimulation block change (−10 to +10 tri-als) or a corresponding block change without a preceding micro-stimulation (non-stimulation).

To assess within session changes in LFP amplitude this procedure was repeated for all block changes (stimulation or non-stimulation) and two frequency by trial matrices were obtained per ses-sion by averaging across all stimulated and non-stimulated block changes respectively. To test for statistical differences between activity during stimulated and non-stimulated block changes the stimulation matrix was contrasted against the non-stimulated matrix to generate a difference matrix for each session. A one-sample t-test was applied to each time-frequency bin of this group differ-ence matrix to test the hypothesis that there was no difference between the stimulation and non-stimulation conditions. The resulting statistical spectra were thresholded at z = 1.66 and cluster cor-rected at p = 0.05 with 1000 permutations used for the cluster correction). This procedure was re-peated separately for both the high-frequency micro-stimulation and control micro-stimulation pro-tocols.

A further analysis assessing changes in LFP amplitude across sessions was carried out on all three session types (control non-stimulation sessions, high-frequency stimulation sessions and control stimulation sessions), as with the across session behavioural analysis (see above) LFP amplitudes from the stimulation sessions were split into stimulated and non-stimulated blocks yielding five conditions. In addition, LFP amplitudes were split by trial type into four groups, explorative trials (rule errors in the final 10 trials of a block) and pre-block change correct trials, perseverative errors (rule error in the final 10 trials of a block) and post-block change correct trials. Differences between trial types in each of the five stimulation conditions were assessed using one sample t-tests on the difference between the groups (e.g., the difference between beta amplitude in non-stimulation con-trol sessions and stimulated blocks in high-frequency stimulated sessions). P-values were corrected for multiple comparisons using Holm-Bonferroni correction.

## Acknowledgments

We thank Biomedical Services staff (providing expert veterinary, and high standards of animal technician support), and recent lab members Martin O’Neill, Erica Boschin and Zhemeng Wu for their assistance and technical support.

## References

1. Tisserand, D.J., et al., Regional frontal cortical volumes decrease differentially in aging: an MRI study to compare volumetric approaches and voxel-based morphometry. Neuroimage, 2002. 17(2): p. 657–69.

2. Mansouri, F.A., et al., Managing competing goals - a key role for the frontopolar cortex. Nat Rev Neurosci, 2017. 18(11): p. 645–657.

3. Braver, T.S. and S.R. Bongiolatti, The role of frontopolar cortex in subgoal processing during working memory. Neuroimage, 2002. 15(3): p. 523–36.

4. Boorman, E.D., et al., How green is the grass on the other side? Frontopolar cortex and the evidence in favor of alternative courses of action. Neuron, 2009. 62(5): p. 733–43.

5. Koechlin, E., et al., The role of the anterior prefrontal cortex in human cognition. Nature, 1999. 399(6732): p. 148–51.

6. Koechlin, E., C. Ody, and F. Kouneiher, The architecture of cognitive control in the human prefrontal cortex. Science, 2003. 302(5648): p. 1181–5.

7. Mansouri, F.A., et al., Behavioral consequences of selective damage to frontal pole and posterior cingulate cortices. Proc Natl Acad Sci U S A, 2015. 112(29): p. E3940–9.

8. Boschin, E.A., C. Piekema, and M.J. Buckley, Essential functions of primate frontopolar cortex in cognition. Proc Natl Acad Sci U S A, 2015. 112(9): p. E1020–7.

9. Mansouri, F.A., M.J. Buckley, and K. Tanaka, Mapping causal links between prefrontal cortical regions andintra-individual behavioral variability. Nature Communications, 2024. 15(1): p. 140.

10. Feizpour, A., et al., The role of frontopolar cortex in adjusting the balance between response execution and action inhibition in anthropoids. Progress in Neurobiology, 2024. 241: p. 102671.

11. Dreher, J.-C., et al., Damage to the Fronto-Polar Cortex Is Associated with Impaired Multitasking. PLOS ONE, 2008. 3(9): p. e3227.

12. Rowe, J.B., et al., Is the prefrontal cortex necessary for establishing cognitive sets? J Neurosci, 2007. 27(48): p. 13303–10.

13. Volle, E., et al., The role of rostral prefrontal cortex in prospective memory: a voxel-based lesion study. Neuropsychologia, 2011. 49(8): p. 2185–98.

14. Hogeveen, J., et al., What Does the Frontopolar Cortex Contribute to Goal-Directed Cognition and Action? J Neurosci, 2022. 42(45): p. 8508–8513.

15. Tsujimoto, S., A. Genovesio, and S.P. Wise, Neuronal activity during a cued strategy task: comparison of dorsolateral, orbital, and polar prefrontal cortex. J Neurosci, 2012. 32(32): p. 11017–31.

16. Tsujimoto, S., A. Genovesio, and S.P. Wise, Evaluating self-generated decisions in frontal pole cortex of monkeys. Nat Neurosci, 2010. 13(1): p. 120–6.

17. Tsujimoto, S. and A. Genovesio, Firing Variability of Frontal Pole Neurons during a Cued Strategy Task. J Cogn Neurosci, 2017. 29(1): p. 25–36.

18. Ferrucci, L., et al., Social monitoring of actions in the macaque frontopolar cortex. Progress in Neurobiology, 2022. 218: p. 102339.

19. Nougaret, S., et al., Neurons in the monkey frontopolar cortex encode learning stage and goal during a fast learning task. PLOS Biology, 2024. 22(2): p. e3002500.

20. Ainsworth, M., et al., Viewing ambiguous social interactions increases functional connectivity between frontal and temporal nodes of the social brain. J Neurosci, 2021. 41(28): p. 6070–86.

21. Sliwa, J. and W.A. Freiwald, A dedicated network for social interaction processing in the primate brain. Science, 2017. 356(6339): p. 745–749.

22. Buckley, M.J., et al., Dissociable components of rule-guided behavior depend on distinct medial and prefrontal regions. Science, 2009. 325(5936): p. 52–8.

23. Altmann, E.M., Advance Preparation in Task Switching: What Work Is Being Done? Psychological Science, 2004. 15(9): p. 616–622.

24. Fouragnan, E.F., et al., The macaque anterior cingulate cortex translates counterfactual choice value into actual behavioral change. Nat Neurosci, 2019. 22(5): p. 797–808.

25. Daw, N.D., Advanced Reinforcement Learning, in Neuroeconomics. 2014. p. 299–320.

26. Lundqvist, M., et al., Gamma and Beta Bursts Underlie Working Memory. Neuron, 2016. 90(1): p. 152–164.

27. Moeller, S., et al., The effect of face patch microstimulation on perception of faces and objects. Nat Neurosci, 2017. 20(5): p. 743–752.

28. Knudsen, E.B. and J.D. Wallis, Closed-Loop Theta Stimulation in the Orbitofrontal Cortex Prevents Reward-Based Learning. Neuron, 2020. 106(3): p. 537–547 e4.

29. Boorman, E.D., T.E. Behrens, and M.F. Rushworth, Counterfactual choice and learning in a neural network centered on human lateral frontopolar cortex. PLoS Biol, 2011. 9(6): p. e1001093.

30. Daw, N.D., et al., Cortical substrates for exploratory decisions in humans. Nature, 2006. 441(7095): p. 876–9.

31. Hogeveen, J., et al., The neurocomputational bases of explore-exploit decision-making. Neuron, 2022. 110(11): p. 1869–1879.e5.

32. Zajkowski, W.K., M. Kossut, and R.C. Wilson, A causal role for right frontopolar cortex in directed, but not random, exploration. eLife, 2017. 6: p. e27430.

33. Lundqvist, M., et al., Gamma and beta bursts during working memory readout suggest roles in its volitional control. Nature Communications, 2018. 9(1): p. 394.

34. Roux, F. and P.J. Uhlhaas, Working memory and neural oscillations: α-γ versus θ-γ codes for distinct WM information? Trends Cogn Sci, 2014. 18(1): p. 16–25.

35. Howard, M.W., et al., Gamma oscillations correlate with working memory load in humans. Cereb Cortex, 2003. 13(12): p. 1369–74.

36. Siegel, M., M.R. Warden, and E.K. Miller, Phase-dependent neuronal coding of objects in short-term memory. Proc Natl Acad Sci U S A, 2009. 106(50): p. 21341–6.

37. Miller, E.K., M. Lundqvist, and A.M. Bastos, Working Memory 2.0. Neuron, 2018. 100(2): p. 463–475.

38. Lee, J.H., M.A. Whittington, and N.J. Kopell, Top-down beta rhythms support selective attention via interlaminar interaction: a model. PLoS Comput Biol, 2013. 9(8): p. e1003164.

39. Buschman, T.J. and E.K. Miller, Top-down versus bottom-up control of attention in the prefrontal and posterior parietal cortices. Science, 2007. 315(5820): p. 1860–2.

40. Stokes, M.G., ’Activity-silent’ working memory in prefrontal cortex: a dynamic coding framework. Trends Cogn Sci, 2015. 19(7): p. 394–405.

41. Neubert, F.X., et al., Comparison of human ventral frontal cortex areas for cognitive control and language with areas in monkey frontal cortex. Neuron, 2014. 81(3): p. 700–13.

42. Gu, B.M., et al., Neural correlates of cognitive inflexibility during task-switching in obsessive-compulsive disorder. Brain, 2008. 131(Pt 1): p. 155–64.

43. den Braber, A., et al., Brain activation during cognitive planning in twins discordant or concordant for obsessive-compulsive symptoms. Brain, 2010. 133(10): p. 3123–40.

44. Bramson, B., et al., Anxious individuals shift emotion control from lateral frontal pole to dorsolateral prefrontal cortex. Nature Communications, 2023. 14(1): p. 4880.

45. Schulz, K.P., et al., Response Inhibition in Adolescents Diagnosed With Attention Deficit Hyperactivity Disorder During Childhood: An Event-Related fMRI Study. American Journal of Psychiatry, 2004. 161(9): p. 1650–1657.

46. Arai, S., et al., Altered frontal pole development affects self-generated spatial working memory in ADHD. Brain and Development, 2016. 38(5): p. 471–480.

47. Snelleksz, M., et al., Evidence that the frontal pole has a significant role in the pathophysiology of schizophrenia. Psychiatry Res, 2022. 317: p. 114850.

48. Ainsworth, M., et al., Frontopolar cortex shapes brain network structure across prefrontal and posterior cingulate cortex. Progress in Neurobiology, 2022. 217: p. 102314.

49. Petrides, M. and D.N. Pandya, Efferent association pathways from the rostral prefrontal cortex in the macaque monkey. J Neurosci, 2007. 27(43): p. 11573–86.

50. Mansouri, F.A., M.J. Buckley, and K. Tanaka, Mnemonic Function of the Dorsolateral Prefrontal Cortex in Conflict-Induced Behavioral Adjustment. Science, 2007. 318(5852): p. 987–990.

51. Oostenveld, R., et al., FieldTrip: Open source software for advanced analysis of MEG, EEG, and invasive electrophysiological data. Comput Intell Neurosci, 2011. 2011: p. 156869.

52. Ciranka, S., et al., Asymmetric reinforcement learning facilitates human inference of transitive relations. Nature Human Behaviour, 2022. 6(4): p. 555–564.

53. Mcfadden, D., Conditional logit analysis of qualitative choice behavior. Frontiers in Econometrics, 1973: p. 105–142.

54. Cohen, M.X., A better way to define and describe Morlet wavelets for time-frequency analysis. Neuroimage, 2019. 199: p. 81–86.

