## Supplementary Information for "Frontopolar cortex stimulation induces prolonged disruption to counterfactual processing: insights from altered local field potentials"

**This file includes:**

Materials and Methods

Figs. S1 to S10

References

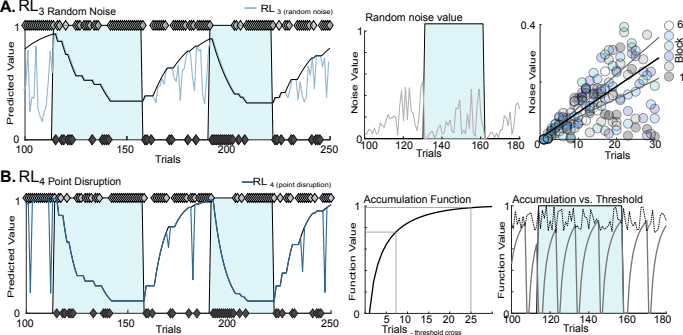
**Supplementary Figures**

**Fig S1 Additional noise driven RL models. A.** RL_3 Random noise_ , values for shape and colour predicted by RL_3_ were modified by addition of random noise to reflect decreased ability to remember the correct rule. Random noise increased in strength during the block. Example of the values of colour predicted by RL_3_ (blue) after modification with randomly generated noise (*left*) with values obtained by RL1 shown for comparison (*black*). Correct trials denoted by light grey diamonds. Examples of the random noise component shown for 80 trials and all blocks of a single session (*right*). Mean random noise value ±SEM denoted with black and grey respectively **B.** RL_4 point disruption_, values for shape and colour switched for a single trial to reflect animals momentarily assigning value on a single trial. Example of the values for colour obtained from a single session by RL_3_ shown in blue with values for RL_1_ in black (*left*). The probability of the values assigned to colour and shape switching increased after a block change until crossing a pseudo random threshold (*right*, see Materials and Methods). The upper and lower bounds of the threshold, and the accumulation function are shown in grey and black respectively. Example of the pseudo-random threshold (black) and accumulation function (grey) shown for 80 trials.

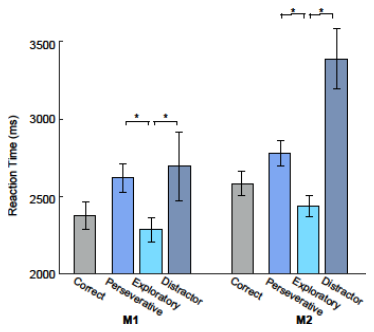

**Fig S2 Reaction time for WCST trial outcomes.** Bar charts showing the reaction times for monkey 1 & 2 for correct (grey) trials, as well as the three possible error outcomes. Rule errors (where the animals chose the incorrect rule) we split into two categories “perseverative errors” made by the animals over the first 7 trials of the block (blue) and “exploratory errors” made over the final 7 trials of the block (cyan). Distractor errors were made when the animals chose the target which didn’t correspond to either rule shown. (blue grey). Significant differences between the mean reaction times were tested with posthoc Bonferroni corrected t-tests.

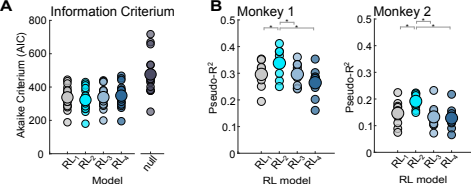

**Fig S3 Addition RL model fit. A.** Akaike Information Criterium (AIC) used to calculated Mcfadden’s pseudo r^2^ shown for all four RL models against the null hypothesis, that animals chose randomly between the stimuli on each trial. **B.** Analysis of the pseudo r^2^ values obtained for all four RL models shown for both M^1^ and M^2^ separately. RL2 chosen + unchosen (light blue) provided a significantly better fit than the other three models in both animals. Asterisk denotes significant differences between model fit at p<0.05)

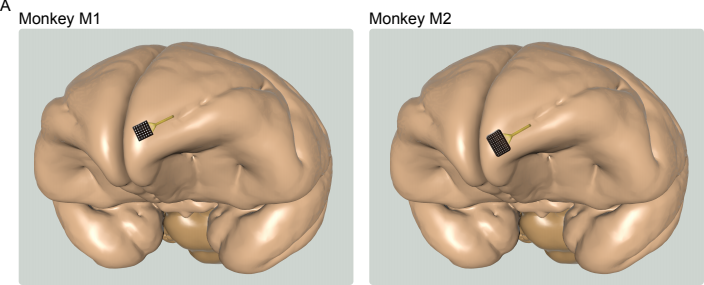

**Fig S4 Electrode locations in M1 and M2. A.** Diagrams showing the location of FPC arrays in M1 and M2 confirmed by post-mortem examination of both animals’ brains. Note difference in array size (32 channel and 64 channel arrays for M1 and M2 respectively).

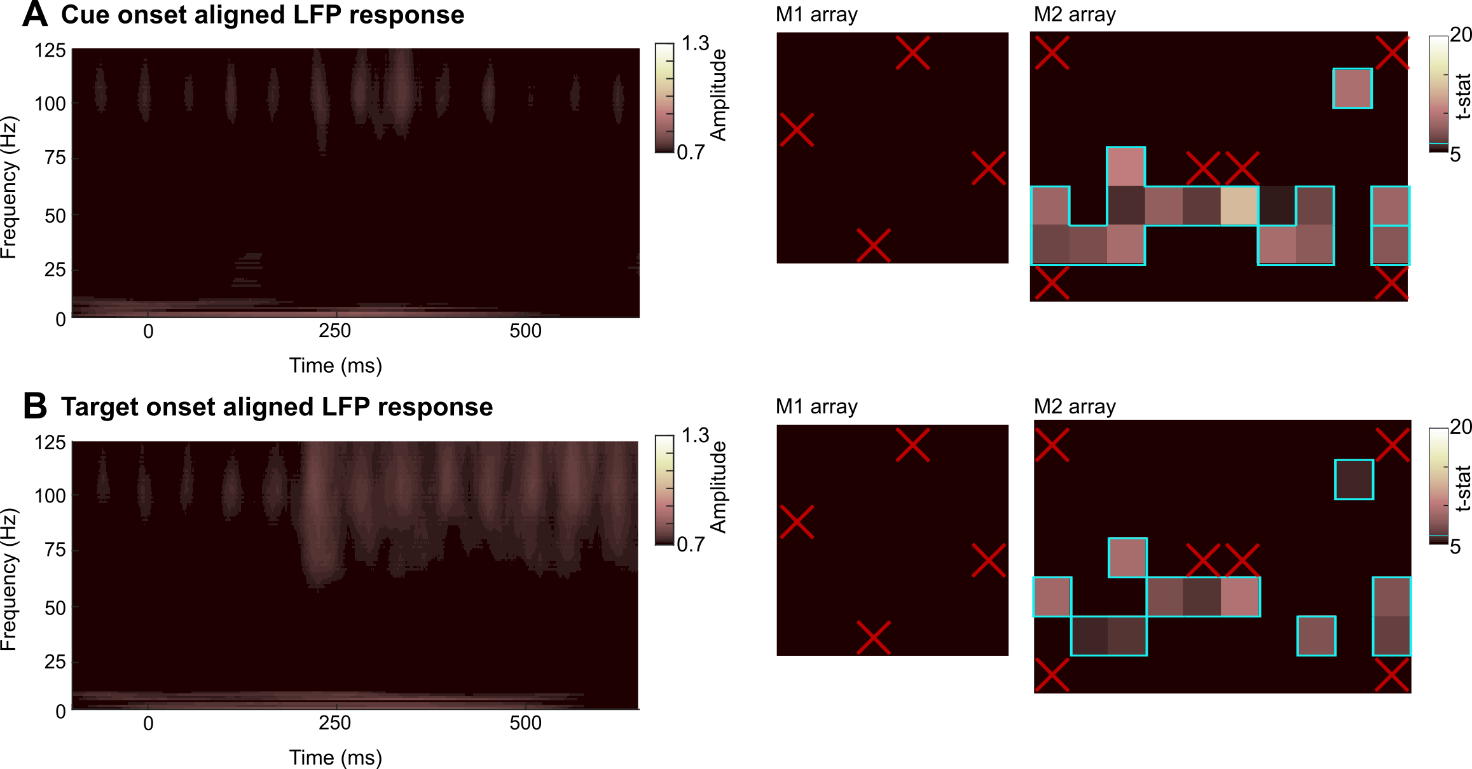

**Fig S5 Field potential activity in FPC is not modulated by onset of cue or target stimuli.** Mean spectrograms calculated from electrodes in FPC showing the absence of gamma activity (*left panel*) and maps showing the limited response in gamma activity observed in arrays implanted in M1 & M2 (*right panels*) aligned to **A.** Cue and **B.** Target onset. Electrodes with significant gamma responses following the relevant trigger outlined in blue. Red crosses indicate reference electrodes

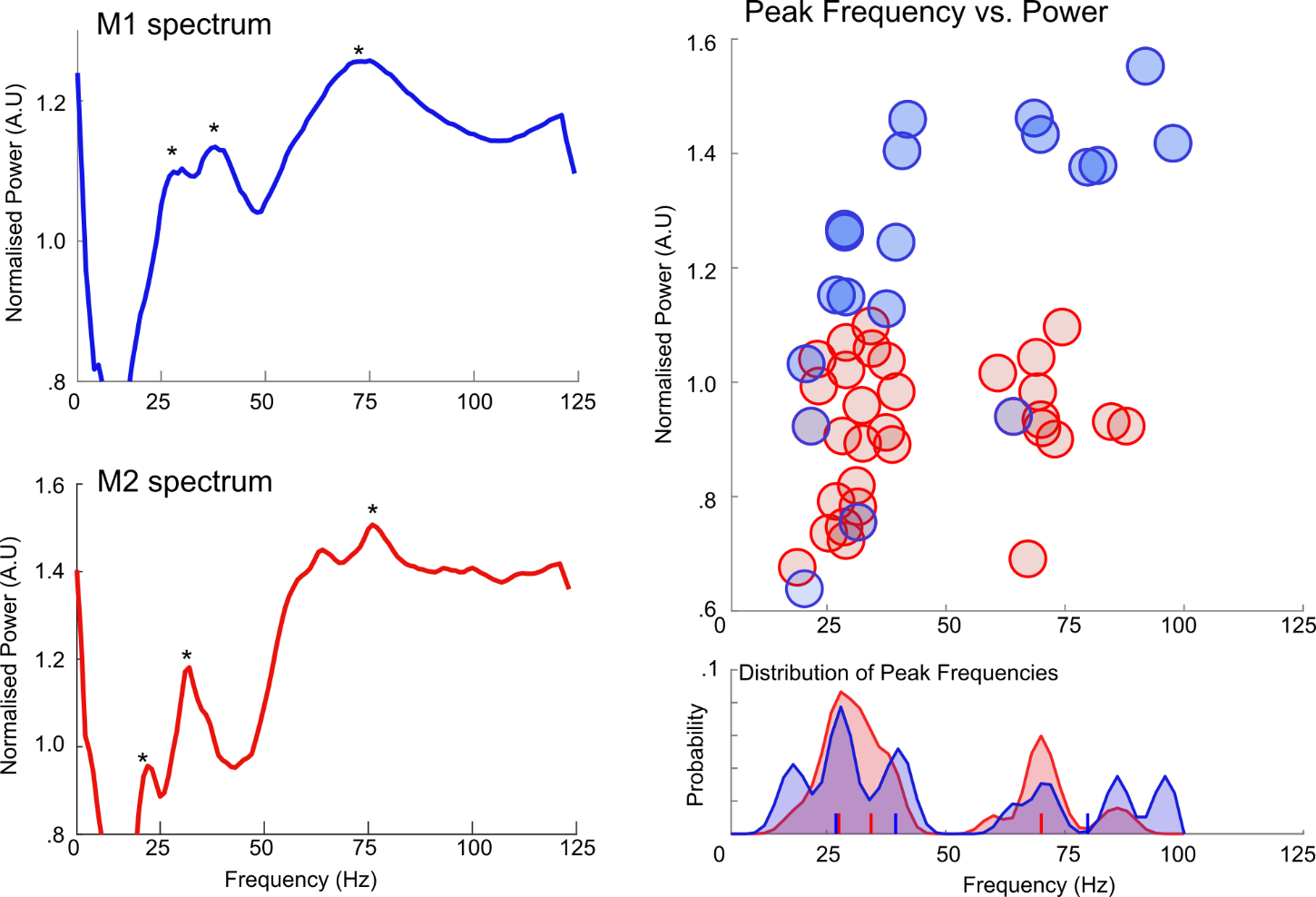

**Fig S6 Overlapping frequencies of beta and gamma activity in M1 and M2. A.** Example choice aligned spectra from M1 (blue) and M2 (red) showing peaks in the beta (15-29 Hz) low gamma (31-45 Hz) and high gamma (55-100 Hz) frequency bands. **B.** Scatter plot showing the peak frequency and amplitude detected in the beta, low and high gamma frequency bands for both M1 and M2. **C.** Summary of the peaks detected for both M1 and M2. Vertical lines denote the mean frequency of peaks in the beta, low and high gamma bands for M1 (blue) and M2 (red) respectively. Note the overlap of peak frequency of activity observed in both animals.

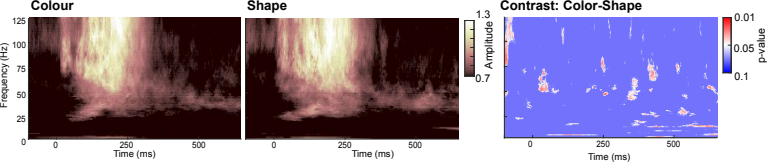

**Fig S7 Choice aligned spectrograms by abstract rule.** Choice aligned spectrograms of LFP activity recorded from FPC split by block (colour and shape) and average across all recording sessions from both animals (left and middle). Statistical spectrogram showing the difference of colour - shape. No significant differences survived cluster correction.

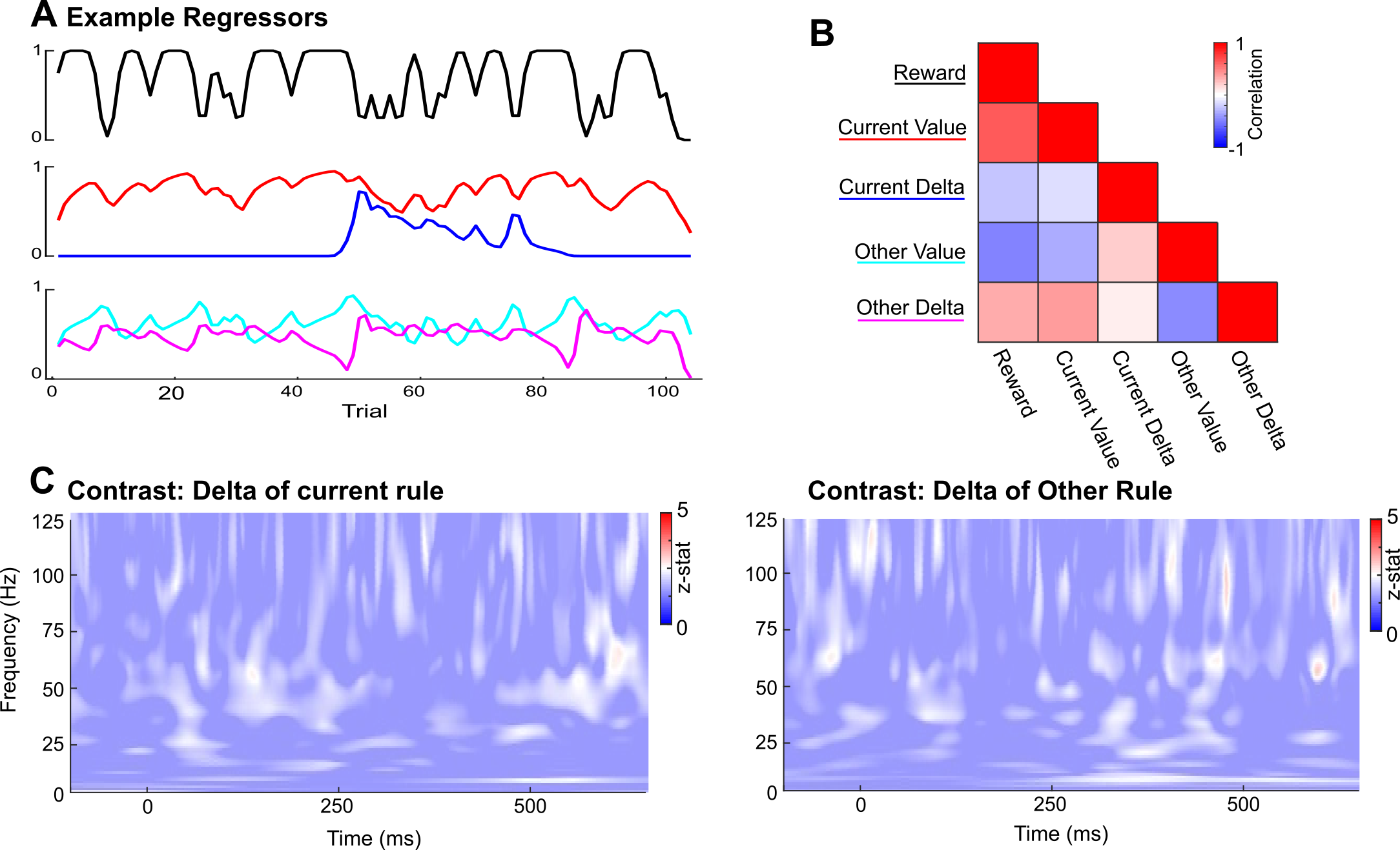

**Fig S8 GLM analysis of choice aligned activity in FPC A.** Example regressors from derived from a single behavioural session input to the GLM analysis. Regressors include reward (black), the estimated value of the current (red) and the other abstract rule (cyan) and the trial-by-trial difference in both values, current rule delta (blue) and other rule delta (purple) **B.** The mean correlation between all five regressors averaged over all sessions from both animals. **C.** Spectrograms showing the Results from the GLM for two contrasts: the delta of the current, and the delta of the other rules.

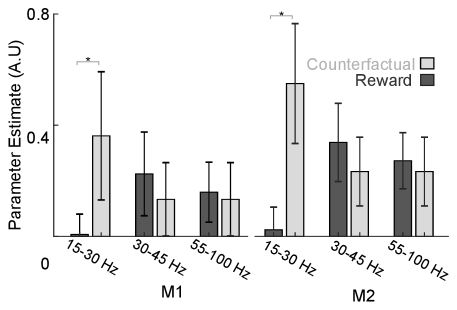

**Fig S9 Summary of the relationship between LFP activity in FPC and counterfactual and reward values for M1 and M2.** Mean parameter estimates obtained from GLM analyses for beta (15-30 Hz), low gamma (30-45 Hz) and high gamma activity (55-100 Hz). Parameter estimates for counterfactual (light grey) and reward (dark grey) shown for both M1 and M2 separately.

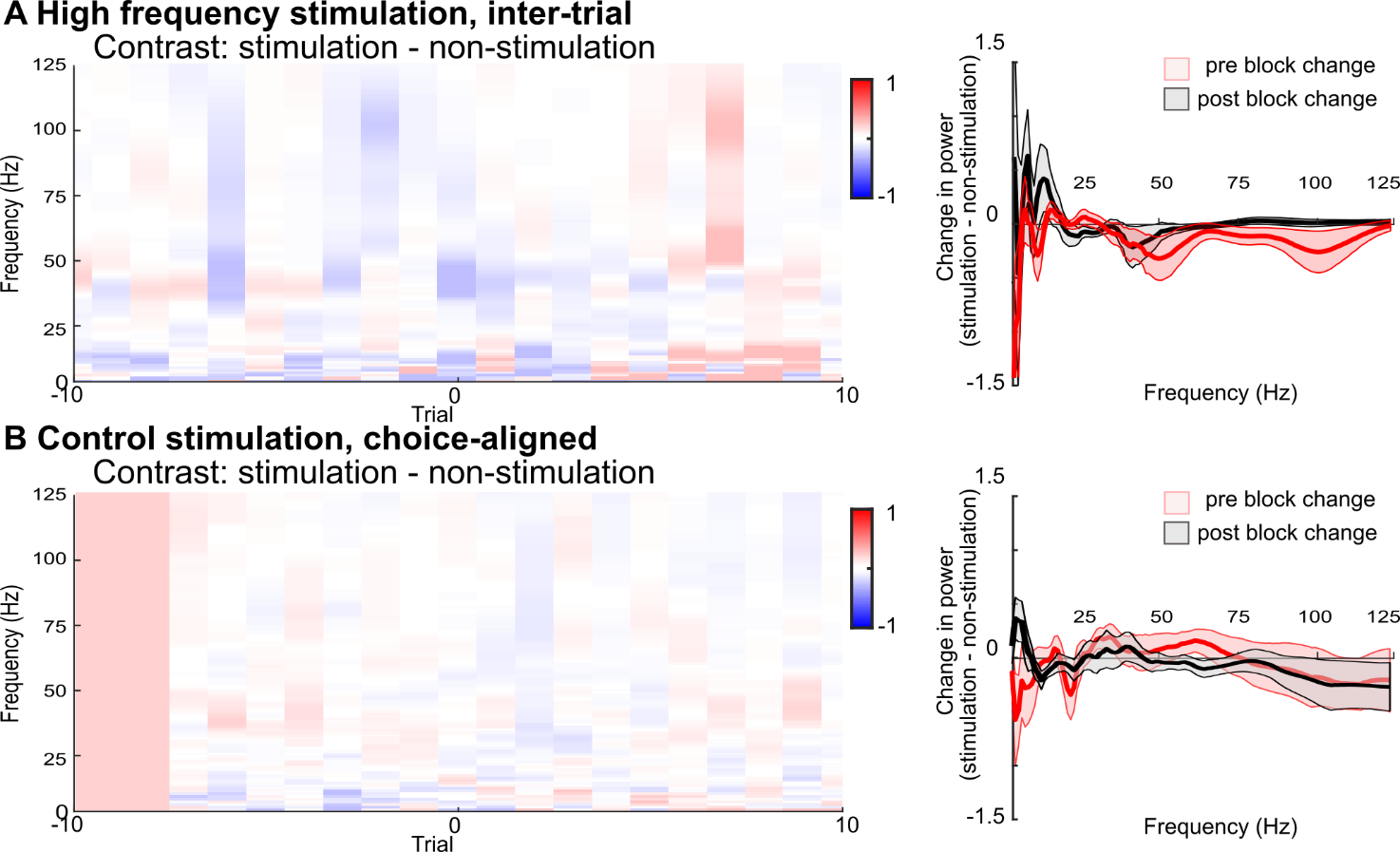

**Fig S10. Trial by-trial spectral analysis showing the difference between the field potential in FPC following** A**)** control stimulation **(B)** non-stimulated blocks within the same sessions. Data shown are 400ms periods of choice aligned **(B)** or inter-trial interval activity **(A)** from the 10 trials preceding and 10 trials following the post-stimulation block change. Significant differences determined using one-sample t-tests (see Materials and Methods), thresholded and cluster corrected at p<0.05. Spectrums showing the average change in LFP power following both stimulation protocols, for the 5 trials pre block change (red) and 7 trials post block change shown on the right (grey). Spectra show mean ±SEM.

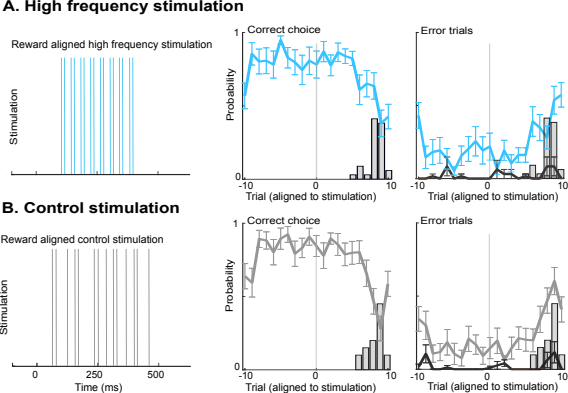

**Fig S11 Stimulation aligned behavioural analysis. A.** Example schematic (*left panel*) showing the high-frequency stimulation protocol (cyan), and the probability of animals performing a correct trial (cyan, *middle panel*) and of making a rule error (cyan) and a distractor error (black, *right panel*) aligned to the stimulated trial for both the high-frequency stimulation. B**.** Example schematic (*left panel*) showing the control frequency stimulation protocol (grey), and the probability of animals performing a correct trial (grey, *middle panel*) and of making a rule error (grey) and a distract or error (black, *right panel*) aligned to the stimulated trial for both the high-frequency stimulation. In all stimulation aligned plots, the probability of the block changing is denoted by grey bars.

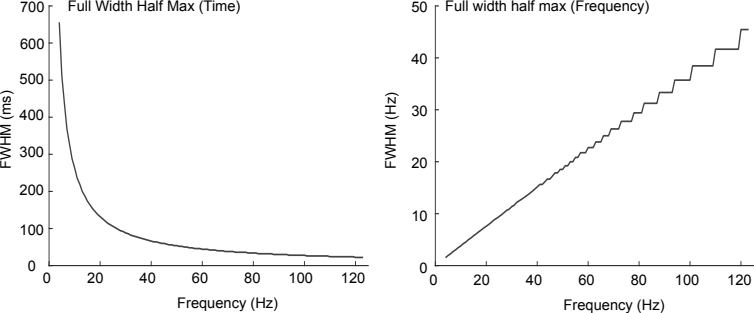

**Fig S12 The full width half max (FWHM) of Morlet wavelets used to decompose LFP activity recorded from FPC.** The FWHM in ms plotted against the central frequency of the wavelets (*left*) and the FWHM converted to frequency domain, again plotted against the central frequency of the wavelet (*right*).

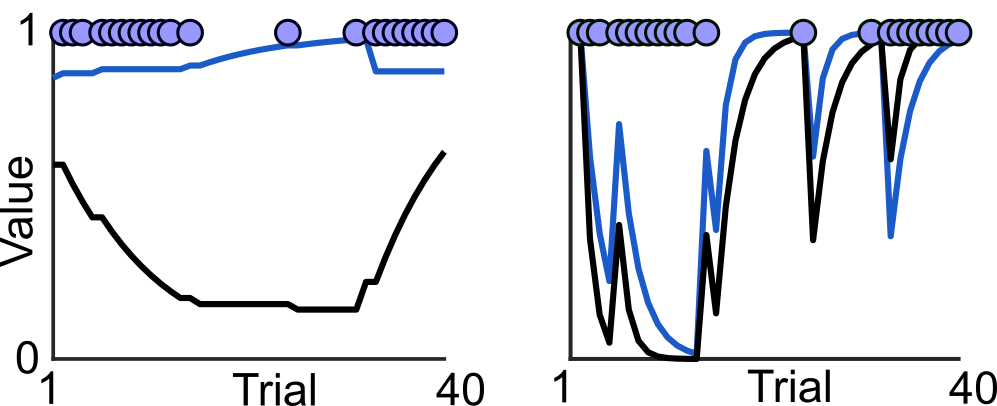

**Fig S13. Comparison of the abstract rule values predicted by RL_1 chosen_ and RL_2 chosen + unchosen._** Plots showing the value of colour (blue) and shape (black) obtained by RL_1 chosen_ (left panel) and RL_2 chosen + unchosen_ (right panel) for 40 examples trials. Blue circles denote trials for which shape was chosen. Note the divergence between the value for shape and colour obtained by RL1 (left) which precludes exploratory behaviour.

**Extended Discussion**

**On the animal's behavioural performance**

Rather than performing the task in an “idealised” form, for example by identifying the correct rule, and exploiting it until a block change, both animals continually made a small number of explorative errors throughout the block. Theoretically an animal could be acting to prolong the rule from changing and hence persist with a currently established ‘cognitive set’ if it made sufficient errors so as never to attain criteria (as criteria was set at 85% correct then the animal would need to ensure its mean error rate was >15% across the session, and that could be achieved either by a sufficient frequency of periodic errors or by short bouts of sustained intentional errors after a good run of corrects. We do not see evidence for either strategy, rather Fig 1C shows the animals continually made rule-errors at a rate which still allowed block changes, and which avoided either extending blocks or decreasing the reward received by the animals. These data suggest the animals may be pre-empting a rule switch, as in this task it was not significantly penalised to prevent the behaviour. It seems likely that such a strategy developed through the animals being sufficiently well trained in the task to have firm concept not only of the two abstract rules (demonstrated by both animals rarely making distractor errors), but also that the counterfactual abstract rule is more likely to become valuable the more reward is gained from the current rule.

This association, between the value attached to the two rules is crucial to the results obtained by our RL modelling of the animals’ choices. RL models which only updated the value of the abstract rule chosen by the animal (model RL_1_) yielded good predictions of the animals’ choices. In particular, and consistent with previous studies [1] these models provided good estimates of the animals’ adaption to changes in abstract rule. However, over the progression of the block the value of the two rules predicted by the model diverged, and don’t reconverge until the next rule switch (Fig S13). Therefore, these models, without additional terms, failed to capture any exploratory behaviour towards the end of blocks. Accurate prediction of these explorative errors required a simple modification of the RL model such that the value of both rules (shape and colour) updated on every trial (RL_2_). In this model both values were updated in the same direction, albeit with the unchosen rule updated at a slower and independently chosen learning rate. Although counter-intuitive, linking the updating of both rules in this fashion both reflects the strategy described above, and significantly improved the RL model fit to the animals’ actual choices.

Indeed, accurate modelling of these explorative errors required modification of our RL models such that the value of both rules were updated in the same direction on every trial (RL_2_). Whilst an optimum strategy may be reasoned to be one in which the correct rule is discerned and then remembered without reference to counterfactual rule until absence of reward indicates time to relearn rule value, the strategy our animals used is arguably a more appropriate one to adopt in volatile probabilistic environments and so unsurprising it is within their repertoire.

**On directed vs. undirected exploration and the inter-species differences in FPC function.**

FPC is a cortical area found in several species of non-human primate. However, there are notable structural and functional differences between the FPC of humans and macaques (the most commonly studied animal model). In humans it has been proposed that a distinction exists between the medial, and lateral sub-division of FPC [2, 3]. With the lateral sub-division reliably activated during tasks requiring directed exploration (e.g., cognitive branching [2, 4, 5]). By contrast the medial sub-division is thought to mediate undirected exploration by allowing assessment of the value of current choices or behaviours. Importantly it has been suggested that the lateral sub-division, and by extension associated cognitive functions (such as cognitive branching) is a feature uniquely associated with humans, with macaque FPC thought to be analogous to the medial sub-division.

The activation of the lateral subdivision of human FPC during cognitive branching is noteworthy as the behaviour requires a more structured consideration of counterfactual task rules (i.e. retaining knowledge of which counterfactual rule to return to). Whilst it remains unclear whether macaques are capable of directed exploration of such a complex nature, our findings provide insight pertinent to the function of FPC in both human and non-human primates. Here the behaviour of our animals suggests their choices, and in particular their adaptation to changes in abstract rule was a form of directed rather than undirected exploration. For example, our animals rarely made distractor errors at any point during testing sessions, and the probability of these errors didn’t increase notably after a change in abstract rule. Further linking macaque FPC with control of directed exploration, microstimulation of FPC caused a significant change in the probability of monkeys making rule errors (either exploratory or perseverative in nature), but no change in the probability of animals making distractor errors.

This link between macaque FPC and directed exploration doesn’t support the proposal that the macaque region can be considered a direct evolutionary ancestor of the medial sub-division of FPC alone. Rather, these findings suggest that macaque FPC supports at least some combined form of the functions associated with both medial and lateral FPC in humans. This interpretation is supported by our demonstration of concurrent signals relating to both assessment of current strategy (reward feedback), and alternative choice value in LFP activity recorded from macaque FPC. By contrast the fMRI evidence from human FPC suggests comparable signals are reliably localised to either the medial or lateral sub-divisions of human FPC [2, 3].

One conclusion based on these findings being that the dissociation of human FPC into medial and lateral sub-divisions represent an evolutionary development whereby functions previously combined in the macaque homologue are processed separately through either the medial or lateral sub-division of FPC. By extension the far greater cognitive complexity and repertoire evident in human explorative behaviour reflects several differences between humans and macaques. Not only the expansion of FPC as a proportion of the brain [6], but also the coexistence of independent processing streams within FPC [2, 3], as well as the innervation of a broader network of regions through the connections of the lateral subdivision [7].

**References**

1.	Steinke, A., F. Lange, and B. Kopp, *Parallel model-based and model-free reinforcement learning for card sorting performance.* Scientific Reports, 2020. **10**(1): p. 15464.

2.	Koechlin, E., et al., *The role of the anterior prefrontal cortex in human cognition.* Nature, 1999. **399**(6732): p. 148-51.

3.	Mansouri, F.A., et al., *Managing competing goals - a key role for the frontopolar cortex.* Nat Rev Neurosci, 2017. **18**(11): p. 645-657.

4.	Dreher, J.-C., et al., *Damage to the Fronto-Polar Cortex Is Associated with Impaired Multitasking.* PLOS ONE, 2008. **3**(9): p. e3227.

5.	Koechlin, E., C. Ody, and F. Kouneiher, *The architecture of cognitive control in the human prefrontal cortex.* Science, 2003. **302**(5648): p. 1181-5.

6.	Semendeferi, K., et al., *Prefrontal cortex in humans and apes: a comparative study of area 10.* Am J Phys Anthropol, 2001. **114**(3): p. 224-41.

7.	Neubert, F.X., et al., *Comparison of human ventral frontal cortex areas for cognitive control and language with areas in monkey frontal cortex.* Neuron, 2014. **81**(3): p. 700-13.
